# Multiscale Analysis of PNPLA2 and PNPLA3 Membrane Targeting

**DOI:** 10.64898/2026.02.12.705593

**Authors:** Amit Kumar, Grace Teskey, Emilio P. Mottillo, Yu-ming M. Huang

## Abstract

Lipid droplets (LDs) are dynamic organelles that regulate cellular lipid storage and mobilization through the coordinated action of LD-associated proteins. Patatin-like phospholipase domain-containing proteins PNPLA2 (ATGL) and PNPLA3 are central regulators of lipid metabolism, yet the molecular mechanisms underlying their membrane targeting and distinct enzymatic activities remain poorly understood. Here, we combine coarse-grained and all-atom molecular dynamics simulations with enhanced sampling to investigate how PNPLA2 and PNPLA3 associate with endoplasmic reticulum (ER) and LD membranes. Despite sharing a conserved N-terminal patatin domain, the two proteins exhibit distinct membrane-binding modes driven by divergent C-terminal amphipathic helices. In both proteins, membrane association is mediated primarily by deep insertion of C-terminal helices, while the patatin domain provides surface contact. PNPLA2 forms a deeply embedded U-shaped helical bundle on LDs that induce pronounced membrane curvature and promote opening of the catalytic dyad, consistent with its high triglyceride lipase activity. In contrast, PNPLA3 engages membranes through a more flexible helical arrangement that maintains a compact catalytic geometry and limits substrate accessibility. Membrane composition further modulates these interactions and leads to protein-specific lipid redistribution and curvature remodeling. Fluorescence microscopy experiments validate the computational predictions and demonstrate that mutation of a single arginine residue within the C-terminal region is sufficient to reduce LD targeting of both proteins. These results establish a mechanistic connection between membrane binding, conformational plasticity, and catalytic regulation in PNPLA2 and PNPLA3. Our work provides molecular insights into how lipid environments tune the function of LD-associated enzymes.

**Author Summary:** LDs are essential cellular organelles that control how fats are stored and released, a process that relies on the precise recruitment and regulation of lipid-metabolizing enzymes. Our work focuses on two closely related enzymes, PNPLA2 (ATGL) and PNPLA3, which play central but distinct roles in lipid metabolism and metabolic diseases. Using a combination of multiscale modeling simulations and fluorescence microscopy, we examine how these proteins recognize and bind to ER and LD membranes. Although PNPLA2 and PNPLA3 share a conserved catalytic core, we show that they interact with membranes in different ways due to differences in their C-terminal amphipathic helices. We find that PNPLA2 forms a deeply embedded helical arrangement that reshapes the membrane and promotes access to its catalytic site, which explains why it typically shows strong lipase activity. In contrast, PNPLA3 adopts a more compact membrane-bound catalytic geometry that limits substrate access and enzymatic activity. We further applied fluorescence microscopy to experimentally validate the computational predictions. The results show that mutation of a single arginine residue within the membrane-binding helix reduces LD targeting. These findings reveal how membrane association and protein conformational dynamics jointly regulate catalytic accessibility and activity.

## Introduction

Lipid droplets (LDs) are intracellular organelles that store neutral lipids and play a central role in cellular energy balance. The patatin-like phospholipase domain-containing proteins PNPLA2 (also known as adipose triglyceride lipase, ATGL) and PNPLA3 are critical regulators of LD turnover and lipid metabolism (1–5). PNPLA2 functions as the primary intracellular triglyceride (TAG) lipase, catalyzing the rate-limiting hydrolysis of TAG in LDs across multiple tissues (6). Its LD localization is critical for this catalytic role and distinguishes it as a key regulator of stored lipid mobilization. Mutations, such as P195L (7), N172K (8), and R221P (9), in PNPLA2 are associated with neutral lipid storage disease with myopathy (NLSDM) (**Table 1**). In contrast, PNPLA3 has minimal basal lipase activity but plays a poorly understood role in hepatic lipid homeostasis (4,10). PNPLA3 localizes to both LD surfaces and endoplasmic reticulum (ER) membranes, which demonstrate its complex regulatory role in lipid metabolism and potentially serving as a bridge between lipid storage and biosynthetic processes (11). PNPLA3 has become a major focus in hepatology, as specific genetic variants significantly influence susceptibility to metabolic dysfunction-associated steatotic liver disease (MASLD) and its progression (**Table 1**) (12–14). This strong clinical association highlights the need to elucidate how PNPLA3 membrane targeting governs its function in health and disease.

**Table 1:** Summary of disease associations, functional roles, and key structural residues of PNPLA2 and PNPLA3.

| Protein | Disease/ Function | Key Residues | References |
| --- | --- | --- | --- |
| **PNPLA2** | neutral lipid storage disease | P195L, N172K, R221P | (7–9) |
|  | catalytic | S47, D166 | (15) |
|  | lipid interaction | I10-G24, T320-P360 | (54) |
|  | ABHD5-mediated activation | N209-N215 | (16) |
| **PNPLA3** | non-alcoholic fatty liver disease | I148M | (51) |
|  | Protective against liver damage with I148M | E434K | (14) |
|  | catalytic | S47, D166 | (15) |
|  | lipid interaction | Q370-V373 | (11) |

No experimentally resolved structures are currently available for PNPLA2 or PNPLA3. However, both proteins share a conserved N-terminal patatin domain (residues 10-180), which contains the catalytic dyad Ser47 and Asp166 essential for hydrolase activity (5,15). In PNPLA2, the C-terminal region (residues 320-360) is critical for LD targeting; experimental studies have shown that removal of this region significantly impairs LD localization (3). Moreover, a recent study identified a short segment (residues N209-N215) as critical for ABHD5-mediated activation of PNPLA2 (16) (**Table 1**), highlighting its role in protein-protein or protein-membrane interactions. In contrast, PNPLA3 has a shorter C-terminal domain with 23 fewer residues compared to PNPLA2. Structural and functional analyses suggest that PNPLA3 relies on a C-terminal segment (residues 345-481) for LD binding, particularly a short QRLV motif (residues 370-373) (**Table 1**). Mutation of this QRLV motif markedly reduces PNPLA3 binding to LDs, which underscores the importance of its C-terminal region in membrane targeting (11).

Although PNPLA2 and PNPLA3 share similar homology, determining how PNPLA2 and PNPLA3 interact with distinct lipid environments is essential for understanding their roles in lipid metabolism and their contributions to metabolic diseases. LDs, composed of a neutral lipid core encased in a phospholipid monolayer, function as dynamic lipid storage organelles. In contrast, the ER membrane, a phospholipid bilayer, is a key site for lipid biosynthesis and protein processing. The atomic-level details of these interactions remain elusive, in part due to the complexity and transient nature of protein-lipid contacts, which are challenging to capture through experimental methods alone.

This study addresses these knowledge gaps by employing a combination of coarse-grained (CG) and all-atom (AA) molecular dynamics (MD) simulations to investigate the membrane-binding mechanisms of PNPLA2 to LDs and PNPLA3 to both LDs and ER membranes. By leveraging multiscale computational approaches, we identified key residues, lipid interactions, and conformational changes that mediate their membrane associations. Our simulations offer a detailed molecular view of the factors governing PNPLA2 and PNPLA3 localization and function, providing new insights into their roles in lipid homeostasis and their potential contributions to metabolic disorders.

## Materials and Methods

### Molecular Models

The amino acid sequences of human PNPLA2 (504 residues) and PNPLA3 (481 residues) were obtained from UniProt (17) (IDs: Q96AD5 and Q9NST1, respectively). A pairwise sequence alignment was conducted using the NCBI BLAST tool (18) to assess their sequence similarity. The initial three-dimensional structures of both proteins were predicted using AlphaFold2 (19). To investigate the membrane interactions of PNPLA2 and PNPLA3, two membrane models were constructed to represent the ER and LD environments. The ER membrane was modeled as a bilayer composed of POPC, DOPE, and SAPI lipids at a molar ratio of 88:37:10. The LD membrane was modeled as a trilayer, consisting of a 4 nm thick core of TAG molecules sandwiched between two ER monolayers (20). The membranes had lateral dimensions of approximately 15 × 15 nm. The LD membrane system included 352 POPC, 148 DOPE, 40 SAPI, and 423 TAG molecules. All systems were solvated with TIP3P water molecules, with a 15 Å thick water layer added above and below the membrane. Potassium and chloride ions were added to achieve a physiological salt concentration of 0.15 M. Coarse-grained models of the protein-membrane systems were first constructed using CHARMM-GUI (21) and PACKMOL (22), and subsequently converted to all-atom models using the Amber tleap program for MD simulations.

### Coarse-grained Molecular Dynamics Simulation

To investigate how PNPLA proteins associate with membranes, we conducted CGMD diffusion simulations. Each protein was initially placed 7.5 nm above the membrane surface in the solution. Simulations were performed using GROMACS version 2020.5 (23) with the MARTINI 3.0 force field (24) applied to proteins, lipids, ions, and solvent. To preserve the secondary and tertiary structure of the protein during simulations, an elastic network model with harmonic restraints was employed. System dimensions and total particle counts for each simulation setup are summarized in **Table S1**.

Energy minimization was executed in two consecutive phases using the steepest descent method (25), each comprising 5000 steps. Equilibration proceeded through five sequential stages, each with decreasing lipid restraint strength and varying integration time steps. Initially, lipid atoms were restrained at 200 kJ/mol/nm² with a 2 fs time step. These restraints were gradually reduced to 100, 50, 20, and 10 kJ/mol/nm² while the integration time step was increased up to 20 fs. No lipid restraints were applied during production runs, which used a constant time step of 20 fs. A full summary of the equilibration protocol is provided in **Table S1**.

Temperature was controlled at 303 K via the velocity rescaling thermostat (26). Semi-isotropic pressure coupling was applied using the Berendsen barostat (27) with a 4 ps coupling constant, a compressibility of 3 × 10⁻⁴ bar⁻¹ in the x and y directions, and a reference pressure of 1 bar. Periodic boundary conditions were used in all three dimensions to mimic bulk membrane behavior. Electrostatic interactions were treated using a relative dielectric constant of 15 and a Coulomb potential smoothly shifted to zero at a cutoff of 1.1 nm. Lennard-Jones interactions were similarly shifted to zero between 0.9 and 1.1 nm. Each system was simulated for 20 µs and repeated in triplicate to ensure statistical reliability.

### Conventional Molecular Dynamics Simulations

Following the membrane-bound conformations obtained from CGMD simulations, we converted the CG systems to all-atom models using the Multiscale Simulation Tool (28). This backmapping procedure places atomistic coordinates around the CG beads based on predefined mapping rules, followed by energy minimization using a modified all-atom force field implemented in OpenMM (29) to resolve unfavorable contacts. All-atom simulations were then prepared using the Amber20 software suite (30). Proteins were assigned parameters from the Amber ff14SB force field (31). Lipid components, including POPC and DOPE, were treated with the Lipid14 force field (32), while parameters for TAG and SAPI were derived from the General Amber Force Field (GAFF) (33). Solvent molecules were modeled with TIP3P water (34), and ionic strength was maintained at 0.15 M KCl using ion parameters developed by Joung and Cheatham (35).

Each system underwent two-stage energy minimization: an initial 2000 steps using the steepest descent algorithm (25), followed by 3000 steps using the conjugate gradient method (36). Systems were gradually heated in two stages with positional restraints applied to lipid atoms. Temperature was first increased from 0 to 100 K over 10 ps, followed by heating to 303 K over 200 ps. This was followed by 10 consecutive 1 ns equilibration runs at 303 K, with all restraints removed. Prior to initiating enhanced sampling protocols, we conducted 50 ns of conventional MD simulations to ensure system stability. All production runs were carried out with a 2 fs integration time step. Temperature was controlled via Langevin dynamics (37) with a collision frequency of 5 ps⁻¹, and pressure was maintained using an anisotropic Berendsen barostat (27). Bond constraints involving hydrogen atoms were enforced using the SHAKE algorithm (38). Long-range electrostatics were treated using the particle mesh Ewald (PME) (39) method with a real-space cutoff of 12 Å.

### Gaussian Accelerated Molecular Dynamics (GaMD) Simulations

GaMD was employed to enhance conformational sampling by adding a harmonic boost potential that flattens energy barriers in the system potential energy landscape (40). This approach improves sampling efficiency without imposing structural restraints (41). Each simulation began with a 50-60 ns preparatory phase, followed by a 1000 ns production run. The preparation phase started with 4-12 ns of standard molecular dynamics at 303 K to collect potential energy profiles. These statistics were used to calculate parameters for the boost potential, including the harmonic force constant and threshold energy. This was followed by a 2-6 ns GaMD equilibration phase with the boost potential applied but without updating the extrema of the potential energy distribution. Subsequently, an additional 50 ns GaMD run was performed with continuous updates to the boost potential. For the final stage, each system underwent a 1000 ns production GaMD simulation using a fixed boost potential. The threshold energy was consistently set to the lower bound (iE = 1), and both dihedral and total potential energies were boosted (igamd = 3). The standard deviation limits for the boost potential on the total energy and dihedral terms were both set to 6.0 kcal/mol (sigma0P = 6, sigma0D = 6). Potential energy statistics were recalculated every 500 ps to ensure consistent boosting parameters. All simulations were carried out in the NPT ensemble using the same thermostat (27), barostat (27), electrostatics (42), and integration parameters as conventional MD. Snapshots were saved every 10 ps for downstream analysis. Three independent GaMD replicates were performed for each system to ensure reproducibility. Full simulation parameters and staging details are provided in **Table S2**.

### Post-MD Analysis

Visual Molecular Dynamics (VMD) (43) and PyMOL (44) were used to visualize simulation systems and create figures and movies. Trajectory analyses were primarily performed using the CPPTRAJ program (45), including calculations of root mean squared fluctuations (RMSF), atom-atom distances, hydrogen bonds, and radius of gyration (Rg). Secondary structure analysis was conducted using PDBsum web server (46), and Ramachandran plots were generated with PROCHECK web server (47). The percentage of contact time between each protein residue and lipid components (POPC, DOPE, SAPI, and TAG) was calculated using MDAnalysis (48). Reported values represent averages from three independent simulation runs. To evaluate how proteins penetrate into membranes, we measured the distance between alpha carbon (Cα) of each residue and the center of mass of nearby membrane phosphorus atoms (within 10 Å in the xy-plane), projected along the z-axis. Membrane surface was analyzed first by calculating the center of mass of each lipid using CPPTRAJ (45), then lipid positions were binned on the xy-plane, and the average height of each bin was computed over the final 400 ns of simulation time.

### Plasmids and Cloning

Human PNPLA2 and PNPLA3 mutants were generated using Overlap Extension PCR using Phusion High-Fidelity DNA Polymerases (ThermoFisher, F530S) as previously described (49). hPNPLA2 WT and hPNPLA3 WT were used as a template. Refer to **Table S3** for primer details. After amplification, the PCR products were digested and cloned in frame into ECFP-N1 or EYFP-N1 using HindIII and AgeI restriction sites. All DNA clones were sequenced and verified for the correct sequence.

### Cell Culture and Transfections

U2OS (ATCC #HTB-96) cells were cultured in DMEM High Glucose (Fisher Scientific, SH30243.01) supplemented with 10% fetal bovine serum (Fisher Scientific, SH3039603) and 5% penicillin/streptomycin (Fisher Scientific, SV30010). Cells were incubated at 37°C and 5% CO2.

U2OS cells were prepared for fluorescent microscopy by seeding on coverslips in a 6 well plate at a density of 300,000 cells/well and allowed to adhere overnight. Cells were transfected using GenJet Plus DNA In Vitro Transfection Reagent (SignaGen Laboratories, #SL100499), with 1µg of total DNA/well and 3µL of GenJet Plus transfection reagent/well. DNA complexes were made in serum-free DMEM. The transfected cells were incubated for 6 hours and the media was removed from the cells and replaced with complete DMEM growth medium with 0.2 mM oleic acid complexed with BSA to promote LD formation. For experiments with PNPLA2, cells were further incubated with 100µM of human ATGL Inhibitor (NG-497, Cayman Chemical Company item# 36886) to further promote LD formation where indicated.

### Fluorescence Microscopy

Cells were transfected with the protein of interest tagged to either ECFP or EYFP. Transfected cells were washed with PBS and fixed with 4% paraformaldehyde. Cells were then stained with either HCS LipidTOX Red neutral lipid stain (Thermo Fisher Scientific, #H34476) or HCS LipidTOX Deep Red neutral lipid stain (Thermo Fisher Scientific # H34477). Diluted 1:1000 in PBS for 30 minutes at room temperature. Images were acquired using a Leica Stellaris confocal microscope at a magnification of 63X. Fluorescent signal from ECFP (excitation 448nm, emission 458nm-523nm), EYFP (excitation 514nm, emission 520nm-543nm), LipidTOX Red (excitation 561nm, emission 589nm-649nm) and LipidTOX Deep Red (excitation 638nm, emission643nm-688nm) was acquired with the Leica LasX software on analog mode. Laser power was set at a threshold that was non-saturating for pixels.

Quantification of fluorescent images was performed by collecting line scan intensities of fluorescence along a linear ROI that included LDs and PNPLA2-ECFP or PNPLA3-EYFP fluorescence in Leica LasX software. The Spearman correlation coefficient between the protein of interest (ECFP or EYFP) and LDs (LipidTOX Red or Deep Red) was calculated based on intensity data. The Spearman correlation coefficient was converted to a z-score using the Fisher’s Z transformation to allow normal distribution of the data. This z-score was weighted by the number of fluorescence intensity data points per image, and a weighted average z-score was generated for each experimental group. To determine significance, the standard error was calculated, and the z-scores were compared on a scale of standard error, generating a z_diff_ statistic. The p-value of the z_diff_ statistic was calculated to determine significance.

## Results

### Sequence Conservation of PNPLA2 and PNPLA3

The sequence alignment of PNPLA2 and PNPLA3 showed moderate conservation, with 46% identity across the full-length proteins. Conservation was higher (54% identity) within the patatin domain (residues 10-180), which harbors the catalytic dyad essential for hydrolase activity. In contrast, the C-terminal regions (residues 181 onward) were more divergent, sharing only 38% identity, consistent with their proposed roles in distinct regulatory or membrane-targeting mechanisms. Pairwise alignment identified key conserved motifs within the patatin domain, including the GXSXG lipase motif (residues 45-49) and the Ser47-Asp166 catalytic dyad (**Figure 1**). The C-terminal regions exhibited lower motif conservation, with PNPLA2 containing additional 23 residues not found in PNPLA3.

**Figure 1:**
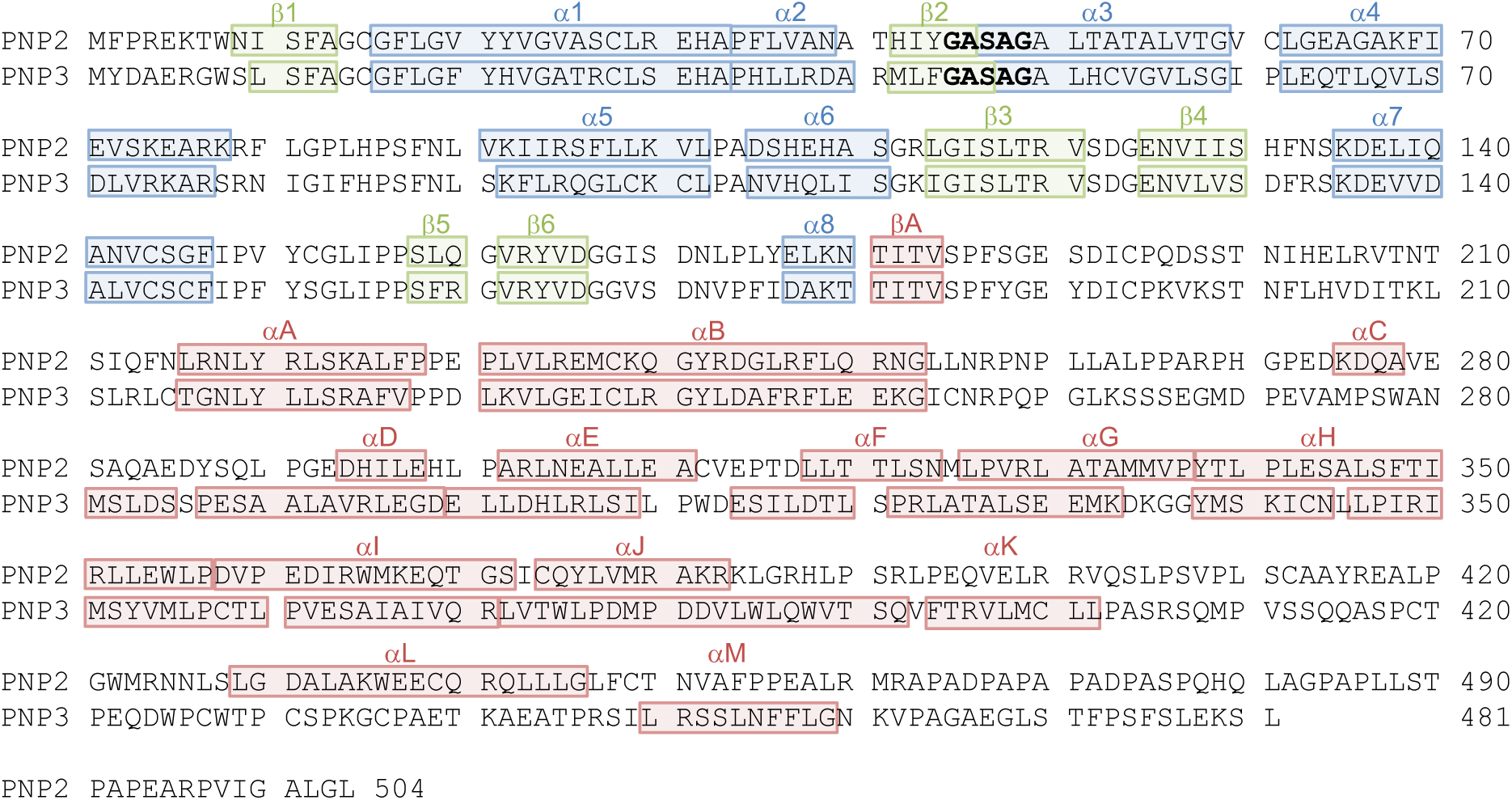
Sequence alignment of PNPLA2 and PNPLA3. Secondary structure elements of the patatin domain are annotated in blue (α-helices) and green (β-strands). Secondary structures in the C-terminal region are shown in red. The conserved GXSXG motifs (residues 45-49) are shown in bold.

The amino acid compositions of PNPLA2 and PNPLA3 reveal similar overall distributions of polar and nonpolar residues, supporting their conserved structural frameworks and enzymatic functions. PNPLA2 consists of 226 polar and 278 nonpolar residues, while PNPLA3 contains 224 polar and 257 nonpolar residues. Both proteins also include similar numbers of positively and negatively charged residues (**Table S4**). This balanced composition supports proper folding and solubility, with hydrophobic cores and surface-exposed polar regions facilitating membrane association. The primary compositional difference lies in the extended C-terminal tail of PNPLA2, which contains 19 additional hydrophobic residues. This hydrophobic extension contributes to distinct membrane-targeting properties of PNPLA2.

### Membrane Interactions of PNPLA2 and PNPLA3

To investigate how PNPLA2 and PNPLA3 interact with membranes at the molecular level, we began by performing CGMD simulations to capture the full-protein diffusion to membranes (**Movie 1-3**). Representative protein-membrane binding poses from CGMD were then converted into all-atom models, which were further equilibrated and analyzed using enhanced GaMD simulations (**Movie 4-6**). To quantify the specific lipid components involved in protein-membrane association, we measured the contact duration between individual residues of PNPLA2/PNPLA3 and various lipid species of ER and LD membranes (**Figure 2**).

**Figure 2:**
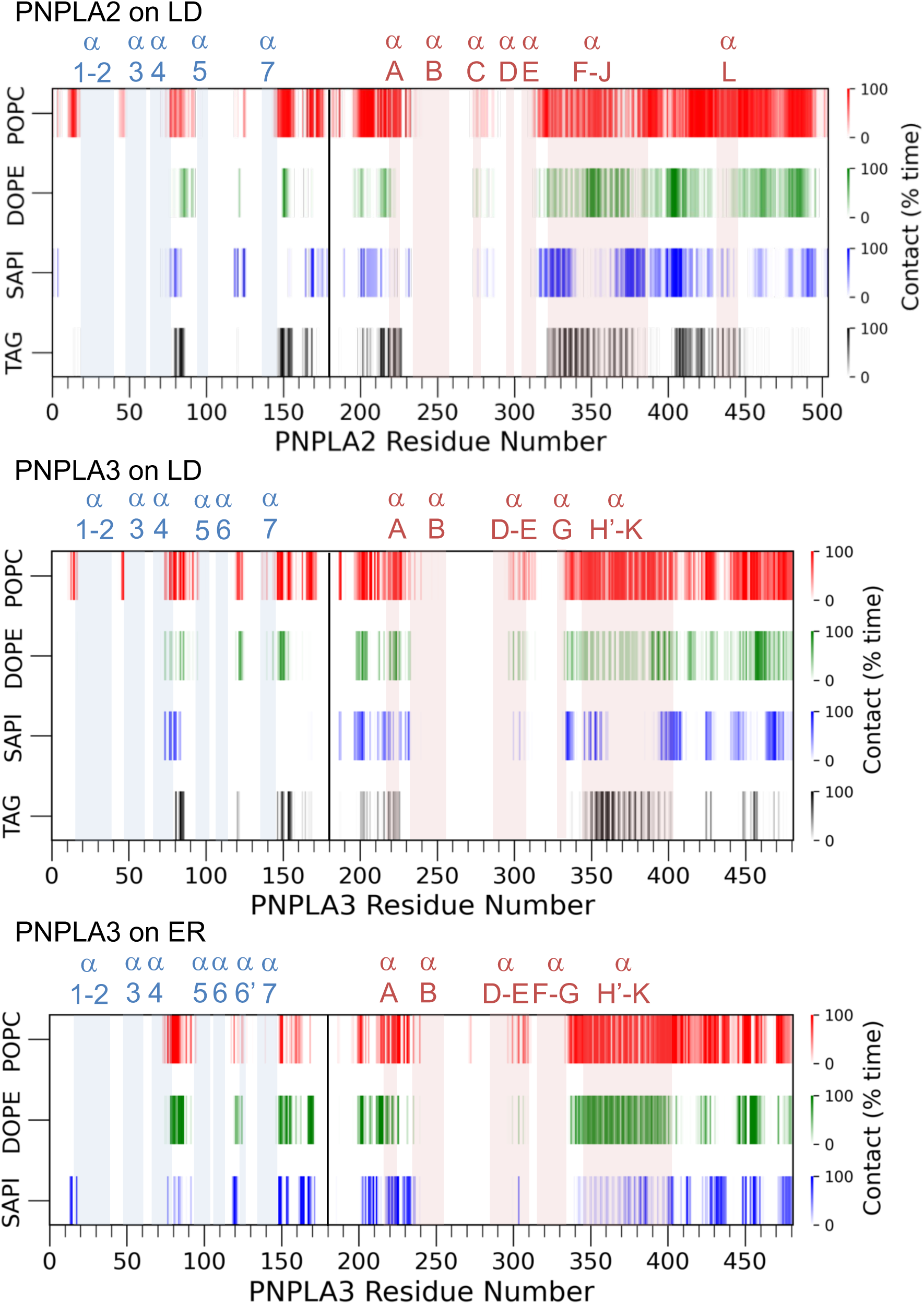
Residue-level contact time between PNPLA proteins and membrane lipids. Contact time (%) between each residue and lipid molecules is shown for PNPLA2 bound to the LD membrane, PNPLA3 bound to the LD membrane, and PNPLA3 bound to the ER membrane. A black vertical line separates the patatin domain (left) from the C-terminal region (right). Key regions that form α-helices are highlighted in blue (patatin domain) and pink (C-terminal region) along the x-axis to indicate their positions relative to the lipid contact patterns. Red, green, blue, and black represent contact time with POPC, DOPE, SAPI, and TAG lipids, respectively. Line color intensity from light to dark reflects increasing contact time from 0% to 100%.

Our results reveal that both the patatin domain and C-terminal region of PNPLA2 and PNPLA3 contribute to membrane interactions. In the patatin domain, membrane association is primarily mediated by surface-exposed loops connecting α-helices in both proteins. In the C-terminal region, both PNPLA2 and PNPLA3 utilize α-helices and terminal loops for membrane binding; however, the specific helices involved differ between the two proteins. PNPLA2 interacts with membranes through helices αF to αJ and forms an additional helix, αL, that engages with the LD surface. In contrast, PNPLA3 primarily uses helices αH′ to αK for membrane binding (**Figure 2 and 3**). PNPLA3 exhibits similar residue-level interactions with both ER and LD membranes.

**Figure 3:**
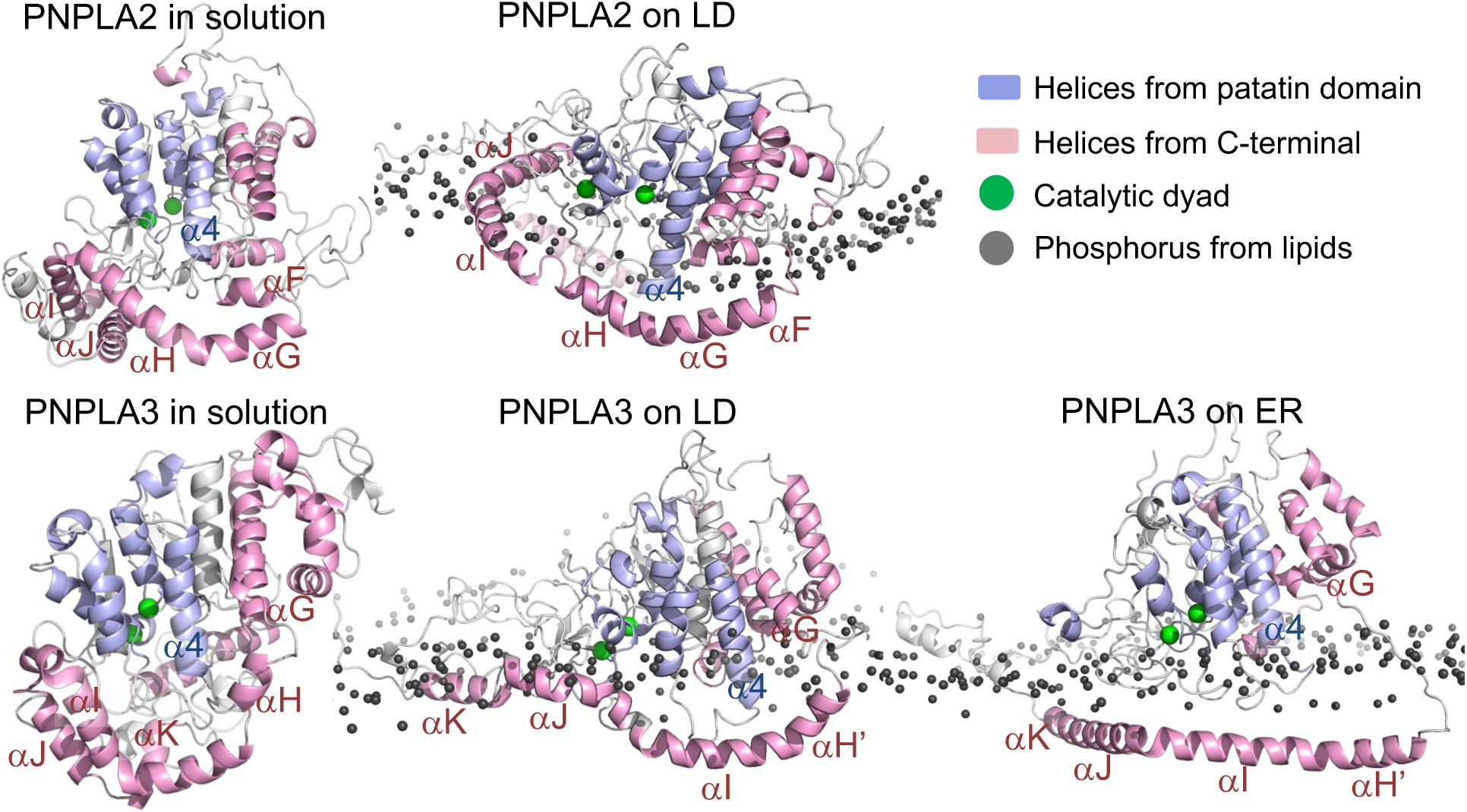
Representative MD snapshots of PNPLA2 and PNPLA3 in different environments. Representative structures from all-atom MD simulations are shown for PNPLA2 and PNPLA3 in three environments: in solution, bound to LD, and bound to ER membrane. Each panel displays the equilibrated conformation of the protein, illustrating environment-dependent structural differences. Blue indicates α-helices from the patatin domain, and pink indicates α-helices from the C-terminal region. The catalytic dyad residues Ser47 and Asp166 are shown as green spheres. Orange spheres represent phosphate atoms from lipid headgroups in the upper leaflet of the membrane, highlighting protein-membrane interaction interfaces.

To assess the extent of membrane penetration by PNPLA2 and PNPLA3, we calculated the vertical distance between each residue and the membrane surface in the ER and LD (**Figure 4A**). A total of 155, 91, and 71 residues are buried within the membrane in the PNPLA2-LD, PNPLA3-ER, and PNPLA3-LD systems, respectively. In all cases, only 9-12 of these residues belong to the patatin domain, while the majority are from the C-terminal region, indicating that membrane insertion is predominantly mediated by the C-terminal. We further quantified the average insertion depth of different protein regions. Across the above three systems, the patatin domain inserts only 2.76 ± 1.89 Å on average, while the C-terminal domain penetrates 5.21 ± 3.62 Å into the membrane, with helices (αG/αH′) reaching depths of 12-16 Å (**Table S5**). Although the specific membrane-binding helices vary among the three systems, deep insertion by C-terminal helices is consistently observed. These data are consistent with mutations in PNPLA2 that lack the C-terminus which impairs LD targeting and causes NLSDM (3). The primary difference between PNPLA2 and PNPLA3 lies in which C-terminal helices interact most deeply with the membrane. In PNPLA2, helices αF to αH penetrate into the membrane interior, while αI and αJ remain closer to the surface (**Figure 3**). In contrast, PNPLA3 shows deep insertion of the entire αH′ to αK region, suggesting more extensive membrane engagement by its C-terminal helices.

**Figure 4:**
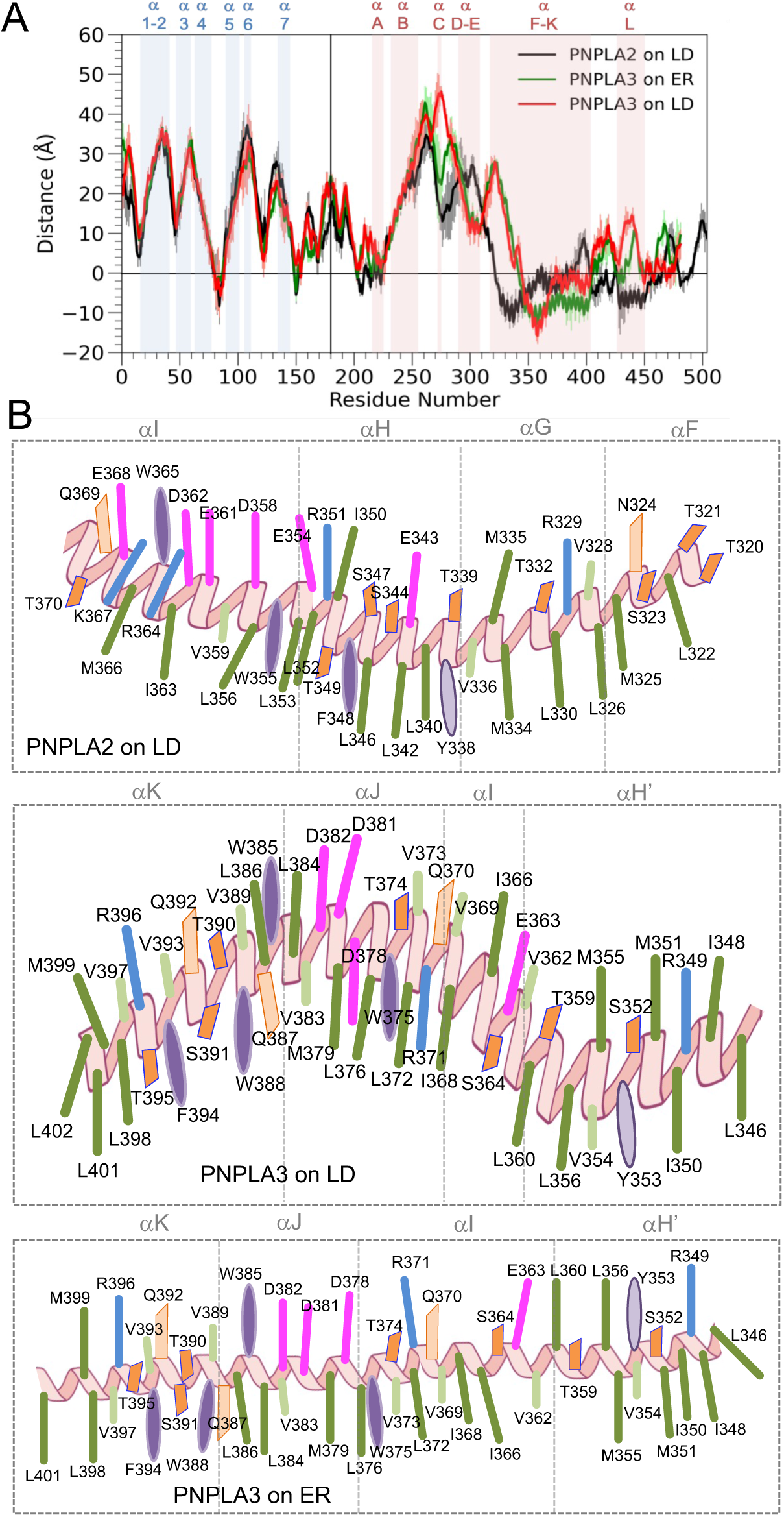
Membrane insertion and amphipathic helix orientation of PNPLA2 and PNPLA3. (A) The average vertical distance (Å) of each residue from the membrane surface in PNPLA2-LD (black), PNPLA3-ER (green), and PNPLA3-LD (red) systems. Negative values indicate membrane insertion. Patatin domain (blue) and C-terminal helices (red) are highlighted, with a vertical black line marking the domain boundary. (B) C-terminal helices from PNPLA2-LD, PNPLA3 in LD- and ER-bound states. Hydrophobic (green and purple), polar (orange), and charged (magenta and blue) residues are labeled.

Both PNPLA2 and PNPLA3 deploy C-terminal amphipathic helices to penetrate the membrane layers, but their lengths, sequences, and insertion depths differ. In PNPLA2-LD complexes, helices αF-αI form a U-shaped bundle: αG and αH present bulky hydrophobic side chains that insert deeply into phospholipid tails and TAG, while polar residues on αF (Ser323) and αI (Arg364) anchor at the phospholipid headgroup layer (**Figure 4B**). In contrast, PNPLA3 on LDs arranges its helices in an inverted U due to polar residues (e.g., Ser364, Arg371, Asp378) on αI and αJ that interact with phospholipid headgroups and restrict insertion depth, whereas αH’ still penetrates deeply via its hydrophobic face (**Figure 4B**). When PNPLA3 associates with the ER membrane, the C-terminal helices retain their amphipathic character but undergo a reorientation. Key polar residues rotate toward the headgroup region while hydrophobic side chains cluster into the bilayer core (**Figure 4B**), which results in a less curved helices along the bilayer surface.

### Mutational Analysis of PNPLA2 and PNPLA3 Validates MD Simulations

To validate the computational models, we performed point mutations targeting residues of PNPLA2 and PNPLA3 identified as important for LD binding. Positively charged arginine side chains typically contact negatively charged phosphates of phospholipid head groups (50). We therefore mutated Arg364 within helix αI of PNPLA2 and Arg349 within helix αH of PNPLA3 to glutamic acid. Both residues were predicted by simulations to be embedded in phospholipids and to contribute directly to membrane association. Wild-type (WT) and mutant proteins were expressed in cells and stained for LDs to enable direct visualization of protein localization relative to LDs.

As shown in **Figure 5A**, WT PNPLA2 localized to LDs and exhibited a clear ring-like enrichment around the LD surface and neutral lipid core (i.e. red, LipidTOX) with minimal cytosolic signal. In contrast, the PNPLA2 R364E mutant showed reduced LD localization accompanied by increased diffuse cytosolic distribution, which indicates impaired LD targeting. Similarly, WT PNPLA3 displayed strong LD association with punctate and ring-like fluorescence patterns at the LD surface. Mutation of Arg349 (R349E) in PNPLA3 resulted in decreased LD enrichment and increased cytosolic staining, consistent with compromised membrane binding (**Figure 5B**). As a quantitative measurement, we performed line scans of PNPLA2 or PNPLA3 signal with LDs (i.e. LipidTox signal), calculating the correlation of signal with LD, comparing WT PNPLA2 and PNPLA3 with their respective mutants. As shown in **Table 2**, the weighted average Z score of WT PNPLA2 line scan correlation with LDs was significantly different compared to PNPLA2 R364E (p = 0.000128). Similarly, WT PNPLA3 line scan correlation with LDs was significantly different compared to PNPLA3 R349E (p=0.0019). These results support the computational models and demonstrate that mutation of a single arginine residue within the C-terminal membrane-binding helices is sufficient to reduce LD targeting of both PNPLA2 and PNPLA3. The findings also highlight the functional importance of specific charged residues embedded within amphipathic helices for stabilizing protein association with LDs.

**Figure 5:**
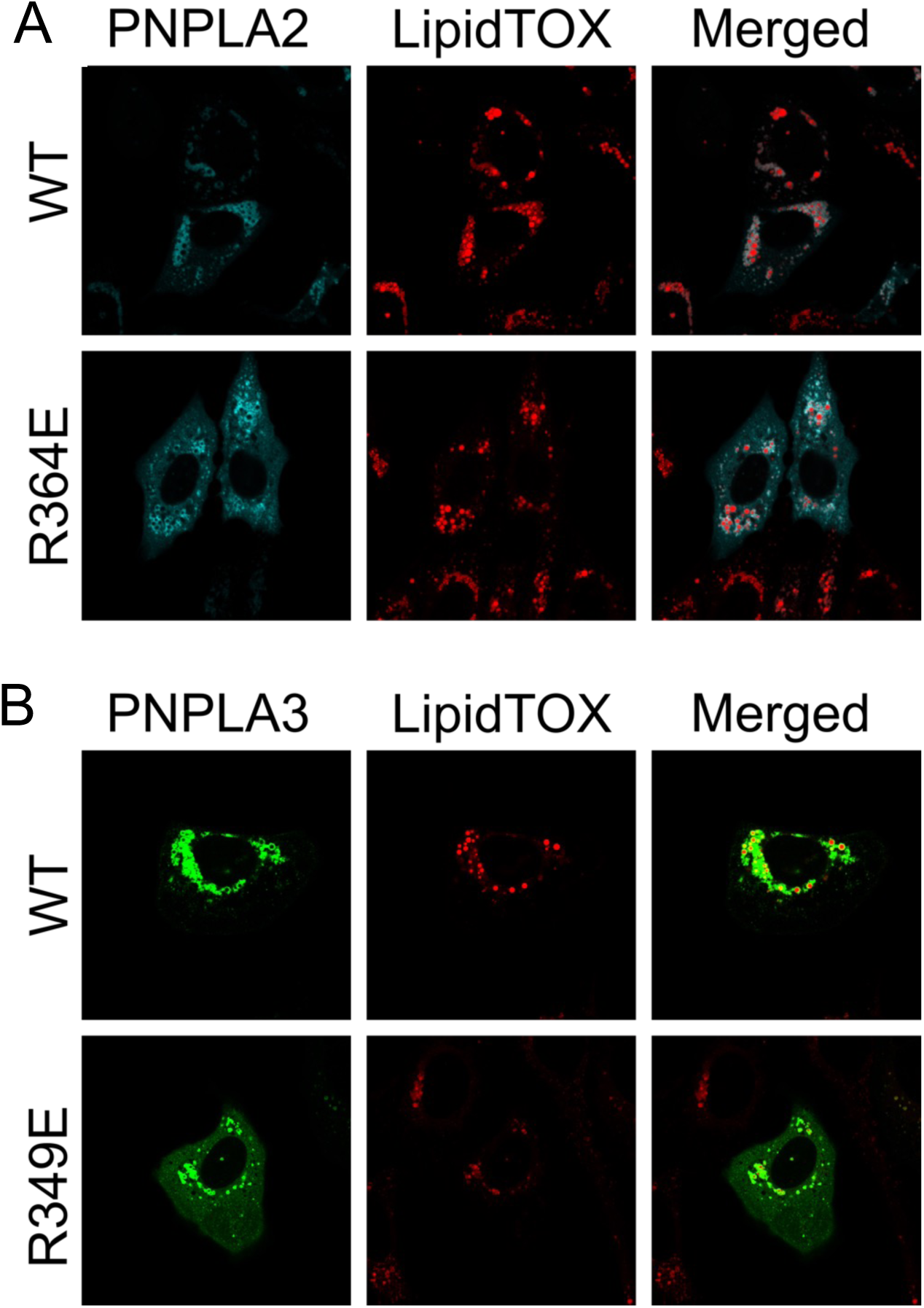
Impact of mutations on PNPLA2 and PNPLA3 LD targeting. (A) Co-localization of WT-PNPLA2-ECFP or PNPLA2 R364E-ECFP (cyan) with LipidTOX Deep Red (red). Cells treated with 0.2 mM oleic acid and 100 µM of human ATGL Inhibitor overnight. (B) Colocalization of WT PNPLA3-EYFP or PNPLA3 R349E-EYFP (green) with LipidTOX Deep Red (red). Cells treated with 0.2 mM oleic acid overnight. Data are representative of three independent experiments performed in technical duplicates for transfections.

**Table 2:** Calculation of weighted line scan correlations of WT PNPLA2 or PNPLA3 and mutants with LDs. PNPLA2 R364E and PNPLA3 R349E correlations were significantly different from their respective WT counterpart.

|  | **Weighted Average Z** | **SE** | **Z<sub>diff</sub>** | **p Value (2-Tailed)** |
| --- | --- | --- | --- | --- |
| **PNPLA2 WT** | 1.005731 | 0.025187 | 3.830832 | 0.000128 |
| **PNPLA2 R364E** | 0.909245 |  |  |  |
| **PNPLA3 WT** | 1.018926 | 0.0252879 | 3.104535 | 0.001900 |
| **PNPLA3 R349E** | 0.940419 |  |  |  |

### Membrane Binding Induces Secondary Structure Changes in PNPLA2 and PNPLA3

We first investigated the secondary structure features of PNPLA2 and PNPLA3 in solution by performing molecular modeling followed by GaMD simulations to equilibrate their structures. The patatin domain of both PNPLA2 and PNPLA3 comprises 8 α-helices and 6 β-strands, organized into a canonical α/β hydrolase fold characteristic of patatin-like phospholipases. This fold features a central five-stranded β-sheet flanked by five major α-helices, with the catalytic triad (Ser47 and Asp166) positioned at the interface (**Figure 3** and **Table S6**). The C-terminal region represents the primary structural divergence between the two proteins. In PNPLA2, this region includes 8 major α-helices and 1 β-strand, while PNPLA3 contains 12 major α-helices and 1 β-strand, indicating a more elaborate helical architecture in PNPLA3. Notably, the C-terminal regions of both proteins are predominantly helical. MD simulations revealed flexibility in the C-terminal regions, with the three replicas showing variations in helical and loop conformations. In PNPLA2, the C-terminal region displayed 12-16 α-helices across trajectories (**Figure S1A**). In PNPLA3, the C-terminal helix count varied between 13 and 20 (**Figure S1B**). Both systems show intermittent loop-to-helix transitions, underscoring the structural dynamics of PNPLA C-terminal region.

Upon membrane binding, the patatin domain retained its overall structural integrity and conserved α/β hydrolase fold across all simulations. However, localized alterations in secondary structure emerged, which reflects subtle adaptations to the membrane environment. For example, α8 converted into a loop conformation following membrane binding in both PNPLA2 and PNPLA3. The C-terminal regions of both proteins, known for their intrinsic flexibility, also exhibited membrane-dependent structural dynamics. Although they retained an overall helical architecture, the number of α-helices varied across simulation replicas depending on the membrane environment. In PNPLA2 bound to LDs, the C-terminal region contained 11-17 α-helices. In PNPLA3, the C-terminal formed 11-14 helices when bound to ER membranes and 13-16 helices when bound to LDs. These results highlight dynamic loop-to-helix transitions and suggest that the C-terminal regions of PNPLA2 and PNPLA3 adapt their structure in response to membrane composition and curvature. Detailed secondary structure analysis is provided in **Table S6** and **Figure S1C, D, E**.

The Ramachandran plot analysis confirmed the stereochemical quality of both solution and membrane-bound models of PNPLA2 and PNPLA3 (**Figure S2**). In solution, 98.8% of residues were located within favored regions, indicating proper backbone dihedral angles and minimal steric clashes. Similarly, the membrane-bound models, PNPLA2-LD (97.4%), PNPLA3-ER (96.7%), and PNPLA3-LD (97.0%), also showed a high percentage of residues in favored regions. These results demonstrate that all models have well-formed backbone conformations and confirm their reliability for MD simulations and subsequent structural analysis.

### Membrane Binding Alters Conformational Flexibility and Compactness of PNPLA2 and PNPLA3

To evaluate structural flexibility, we calculated the root mean square fluctuation (RMSF) for each residue in PNPLA2 and PNPLA3 from three simulation replicas (**Figure 6** and **Table S7**). In solution, the patatin domain of both proteins was highly stable, with average RMSF values of 0.85 Å for PNPLA2 and 0.84 Å for PNPLA3. In contrast, the C-terminal regions were more dynamic, averaging 1.98 Å for PNPLA2 and 1.57 Å for PNPLA3. These fluctuations were most prominent in loop regions, disordered tails, and terminal helices.

**Figure 6:**
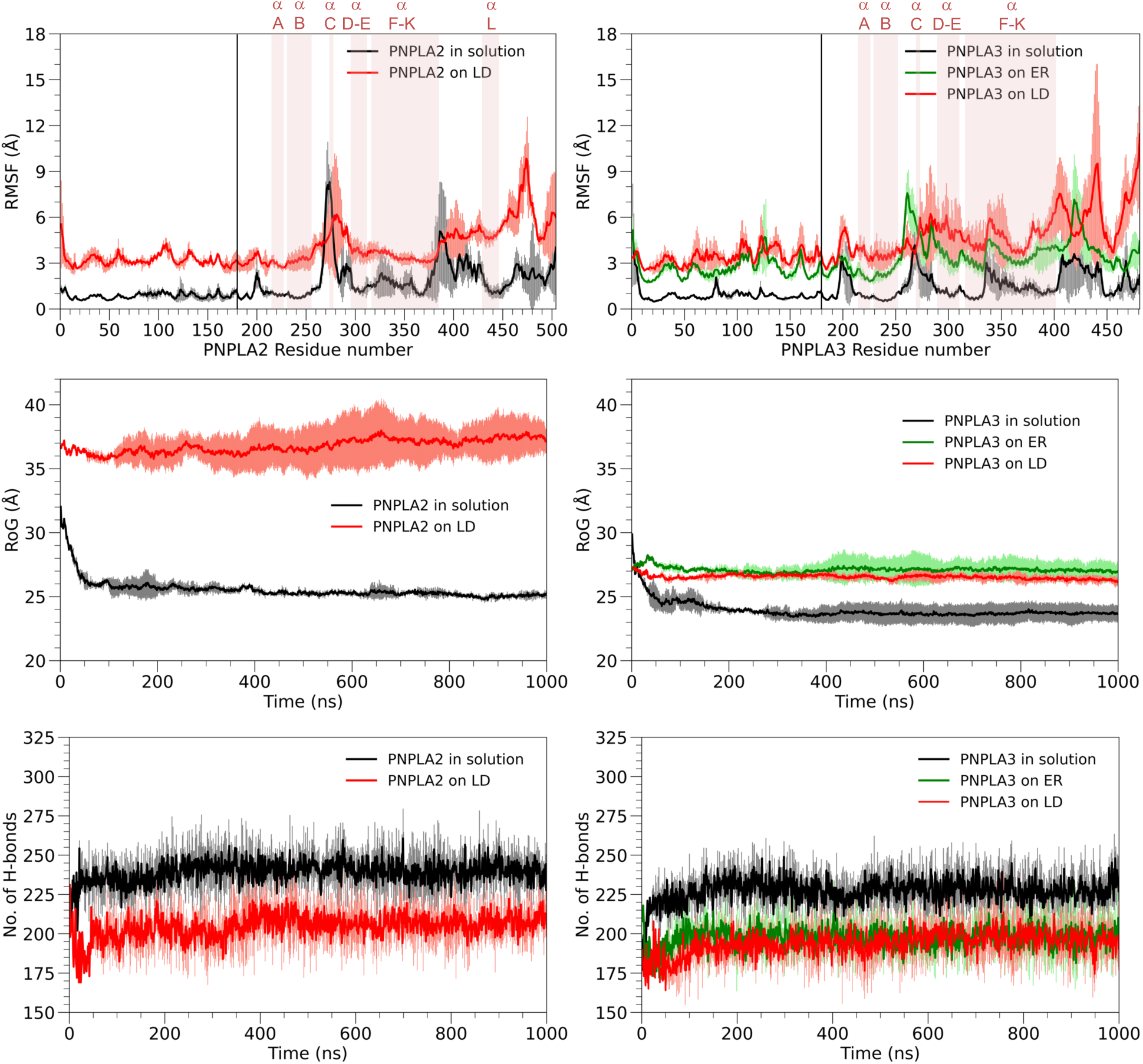
Structural stability of PNPLA2 and PNPLA3 in solution and membrane-bound states. The top row shows residue-wise root-mean-square fluctuation (RMSF) profiles, highlighting atomic flexibility. The middle row presents the radius of gyration (Rg) over simulation time, reflecting global compactness. The bottom row displays the number of intraprotein hydrogen bonds over time, indicating internal stabilization. Left panels correspond to PNPLA2 in solution and bound to LD membranes. Right panels show PNPLA3 in solution and bound to either ER or LD membranes. For detailed numerical values, see Table S6 (RMSF) and Table S8 (Rg and hydrogen bonds).

Membrane binding led to increased fluctuations in both the patatin and C-terminal domains. In LD-bound systems, the patatin domain RMSF rose to 3.18 Å for PNPLA2 and 3.40 Å for PNPLA3. For ER-bound PNPLA3, the value was slightly lower at 2.62 Å with more restrained dynamics. The C-terminal regions became even more flexible, with RMSF values reaching 4.31 Å for PNPLA2-LD, 4.92 Å for PNPLA3-LD, and 3.55 Å for PNPLA3-ER (**Table S7**). The C-terminal helices that interact with the membrane (e.g., αF-αJ in PNPLA2 and αH′-αK in PNPLA3) exhibited elevated fluctuations. For example, the RMSF of helix αH in PNPLA2 increased from 1.59 Å in solution to 3.32 Å upon membrane binding. These enhanced dynamics reflect membrane insertion and increased nonpolar contacts between the protein and lipid tails, enabling flexible yet stable anchoring to the membrane surface.

To evaluate the compactness and internal stability of PNPLA2 and PNPLA3, we analyzed the radius of gyration (Rg) and intraprotein hydrogen bond counts across three independent simulations. In solution, PNPLA2 exhibited an average Rg of 25.52 Å, compared to 23.92 Å for PNPLA3 (**Figure 6 and Table S8**), suggesting a more expanded structure for PNPLA2 and a relatively compact conformation for PNPLA3. Consistently, PNPLA2 formed an average of 239 intraprotein hydrogen bonds, slightly more than the 227 observed for PNPLA3. This difference arises from the longer C-terminal region of PNPLA2, which contributes additional hydrogen-bonding residues and supports protein folding.

Upon membrane association, PNPLA2 adopted a notably more expanded conformation (**Figure S3**). In LD-bound simulations, its Rg increased to 36.85 Å, which reflects conformational rearrangement to accommodate lipid interactions. In contrast, PNPLA3 remained relatively compact, with Rg values of 27.08 Å and 26.52 Å in the ER- and LD-bound states, respectively. These results indicate that membrane binding induces a substantial expansion in PNPLA2, while PNPLA3 maintains a consistent global architecture across environments. Internal hydrogen bonding decreased in both proteins upon membrane binding, due to redistribution of contacts toward membrane lipids. LD-bound PNPLA2 formed an average of 205 intraprotein hydrogen bonds, while ER- and LD-bound PNPLA3 formed 197 and 194 hydrogen bonds, respectively (**Figure 6 and Table S8**). This reduction highlights a structural trade-off in which residues that engage in protein-lipid interactions contribute less to internal stabilization, thereby promoting membrane anchoring and adaptation.

### Lipid Droplet Binding Induces Active Site Opening in PNPLA2

The catalytic dyad consisting of Ser47 and Asp166 is a conserved and functionally essential feature of both PNPLA2 and PNPLA3. Proper spatial positioning of this dyad is critical for enzymatic activity, as it enables nucleophilic attack on lipid substrates. Our simulations revealed that membrane binding and subcellular localization influence the geometry of the catalytic dyad in an environment-specific manner. In PNPLA2, the average distance between Ser47 and Asp166 is 6.20 ± 0.15 Å in solution. Upon LD binding, this distance increases significantly to 11.19 ± 0.45 Å, indicating an open active-site conformation (**Figure 7 and S3**). This structural expansion facilitates lipid substrate access and suggests a catalytically permissive state. In contrast, PNPLA3 exhibits a more restrained dyad geometry. The Ser47-Asp166 distance is 6.52 ± 0.19 Å in solution, increasing slightly to 7.50 ± 0.18 Å in the ER-bound state and remaining similar to 6.33 ± 0.53 Å in the LD-bound state. These conformational differences suggest a potential regulatory mechanism in PNPLA3, in which restricted dyad geometry may limit substrate accessibility. These data are in agreement with and explain the respective greater TAG lipase activity of PNPLA2 vs PNPLA3 (51).

**Figure 7:**
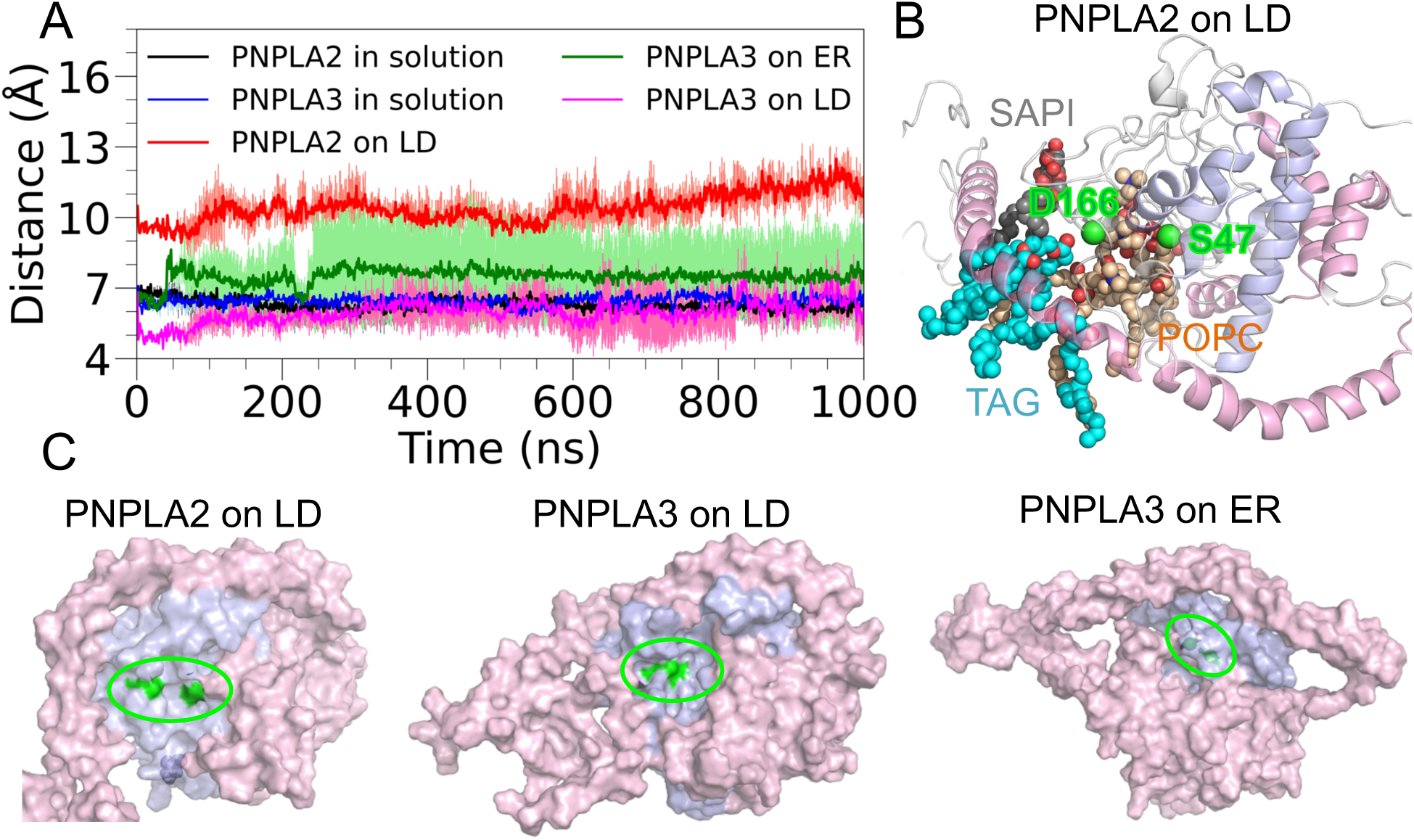
Time evolution of the distance between Ser47 and Asp166 (Cα-Cα) in PNPLA2 and PNPLA3 under different conditions. Distances were calculated over 1 μs GaMD simulations for PNPLA2 in solution (black) and on LDs (red), and for PNPLA3 in solution (blue), on LDs (magenta), and on ER membranes (green). Data represent mean values and standard deviation from three independent simulations.

Analysis of lipid proximity to the catalytic dyad supports this functional divergence. In LD-bound PNPLA2, 7.7 lipid molecules were found within 10 Å of both Ser47 and Asp166, including 2.0 TAG (**Table S9**), indicating active site engagement. In comparison, only 4.27 and 3.56 lipid molecules were observed near the dyad in PNPLA3-LD and PNPLA3-ER, respectively, with TAG almost absent from the PNPLA3 catalytic pocket (**Table S9**). This further suggests reduced substrate accessibility in PNPLA3, despite membrane association. Lipid composition within the catalytic pocket also differed by environment. When bound to LDs, both PNPLA2 and PNPLA3 contained POPC near the catalytic dyad. However, in ER-bound PNPLA3, POPC was absent, and DOPE and SAPI was instead present in the pocket, which indicates membrane composition-specific interactions (**Table S9**).

### Membrane Redistribution and Curvature Change Upon PNPLA2 and PNPLA3 Targeting

Before protein binding, the lipid layers exhibited a relatively homogeneous lateral distribution. Upon PNPLA association, significant changes in lipid spatial distribution were observed. Both PNPLA2 and PNPLA3 disrupted the local lipid environment in a consistent and protein-specific manner. POPC was largely excluded from the membrane area directly beneath the patatin domain of both proteins (**Figure 8**). In contrast, SAPI and TAG accumulated preferentially near the C-terminal regions of both PNPLA2 and PNPLA3. This enrichment is driven by hydrophobic interactions between the lipid tails and the membrane-associated helical segments of the C-terminal domain, which penetrate deeper into the membrane layer. Interestingly, DOPE exhibited divergent distribution patterns. In the presence of PNPLA2, DOPE was excluded from the region under the protein. However, PNPLA3 retained DOPE beneath its patatin domain, particularly when bound to the ER membrane.

**Figure 8:**
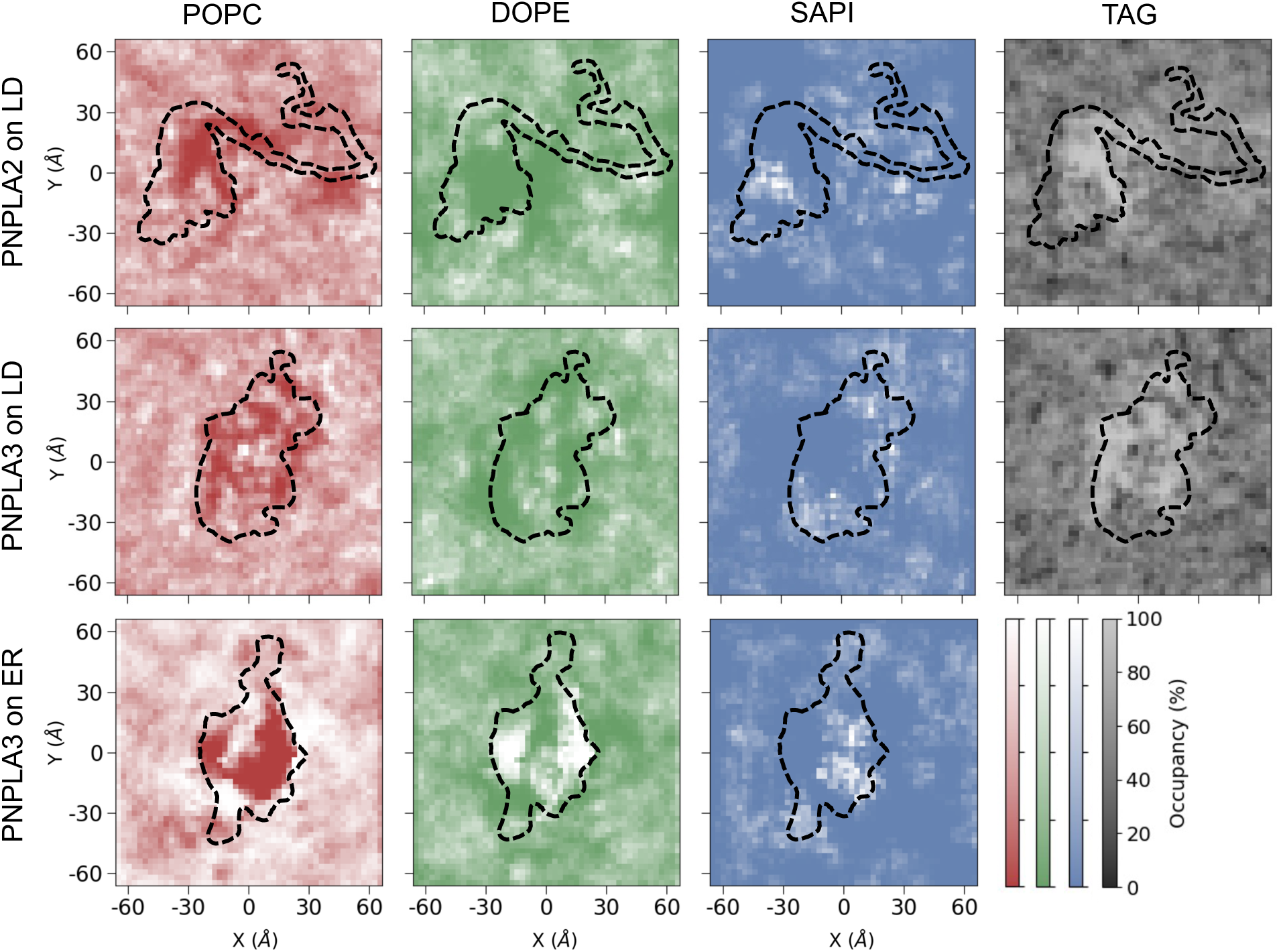
Spatial distribution of lipids around PNPLA2 and PNPLA3 on membrane surfaces. Heatmaps illustrate the distribution of lipids around PNPLA2 and PNPLA3 in different membrane environments. The top, middle, and bottom rows correspond to PNPLA2 bound to the LD, PNPLA3 bound to LD, and PNPLA3 bound to the ER, respectively. POPC, DOPE, SAPI, and TAG lipids are shown in red, green, blue, and gray, respectively. Darker colors indicate regions with low lipid occupancy over the simulation time. Protein locations are outlined with dashed lines for reference.

To evaluate the membrane remodeling induced by PNPLA binding, we quantified the average vertical height of phospholipids and TAG molecules relative to the membrane surface (**Figure 9A and S4**). Using the center of mass of the protein as the origin, lipid heights were radially averaged over defined distances to characterize local curvature (**Figure 9B**). Our analysis revealed that PNPLA2 induces pronounced remodeling of the LD membrane. A distinct negative curvature formed beneath the patatin domain, where phospholipid height was reduced to -9 ± 2.2 Å. In contrast, a positive curvature was observed around the C-terminal region, which shows lipid elevation and upward displacement near this membrane-inserting domain. PNPLA3 also induced local membrane curvature on the LD surface, with a modest depression beneath the patatin domain (-1 ± 2.1 Å), although the magnitude was notably smaller than that caused by PNPLA2. When PNPLA3 bound to the ER membrane, no significant membrane curvature was observed, suggesting that curvature remodeling is membrane-type and protein-specific. In contrast to the lowered phospholipid height beneath the patatin domain, TAG molecules in the PNPLA2-LD system rose to the phospholipid surface, especially around the C-terminal region. This elevation of TAG suggests a potential accumulation or enrichment of neutral lipids driven by the local protein-lipid interactions at the C-terminal helices, which may support lipid recruitment or extraction during enzymatic processing.

**Figure 9:**
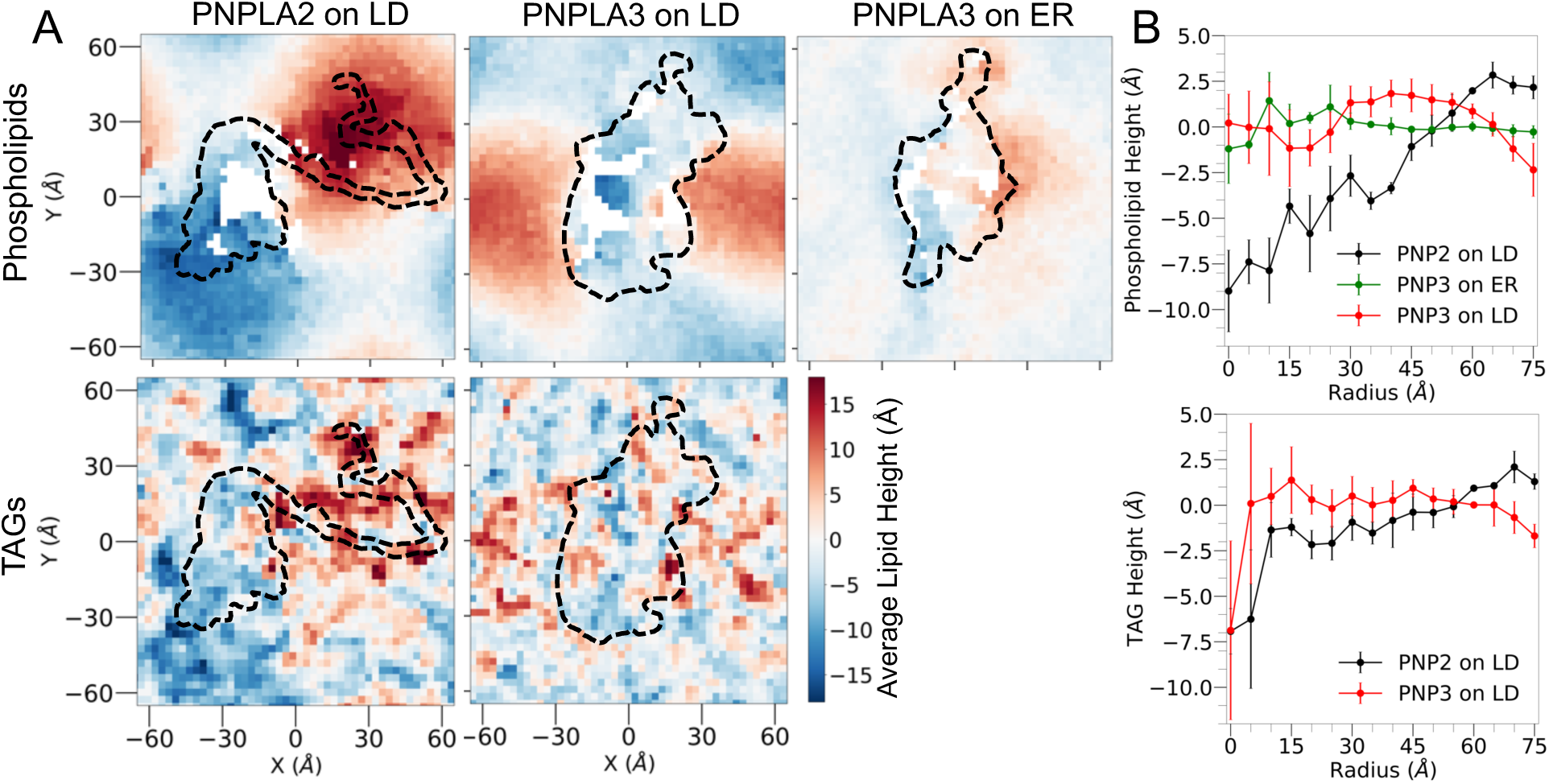
Membrane curvature induced by PNPLA2 and PNPLA3 binding. (A) Height maps of phospholipid and TAG surfaces in the presence of membrane-bound PNPLA proteins, calculated from the last 400 ns of simulation. Columns represent PNPLA2 bound to LD (left), PNPLA3 bound to LD (middle), and PNPLA3 bound to ER (right). The top and bottom rows show the height distribution of phospholipids and TAG, respectively. Heatmaps display lipid height in Å, with color gradients from blue (lower height) to red (higher height). Protein locations are outlined with dashed lines. (B) Radially averaged lipid surface heights as a function of distance from the protein center, computed across three independent simulation replicas. Top and bottom are shown for phospholipids and TAG, respectively. Error bars represent the standard deviation at each radial distance, reflecting variations across three simulation replicas.

## Discussion

This study reveals the molecular basis of membrane binding by two patatin domain-containing lipases, PNPLA2 and PNPLA3, using a multiscale approach. Although these enzymes share high structural homology in their N-terminal patatin domains, our simulations show they adopt distinct membrane-binding modes that may correlate with their divergent functional roles. This dichotomy aligns with prior observations across the PNPLA family (15), where catalytic core residues are conserved despite moderate global sequence identity, supporting the idea that divergence of the C-terminal underlies specialized subcellular localizations and regulatory mechanisms.

Our results reveal that both the patatin and C-terminal domains contribute to PNPLA2 and PNPLA3 membrane association, where the patatin domain provides surface contact while the C-terminal region drives deep membrane insertion, which establishes a dual-factor binding mechanism. This pattern is consistent with other LD-targeting proteins (**Table 3**). For example, alpha beta hydrolase domain containing 5 (ABHD5) interacts with LD by both N-terminal residues (1–43) and a membrane-anchoring domain within the insertion segment (residues 180-230). Another example is the hormone-sensitive lipase (HSL), which employs membrane-targeting through an N-terminal four-helix bundle (residues 1-136) and a C-terminal H-motif (residues 489-538).

**Table 3:**
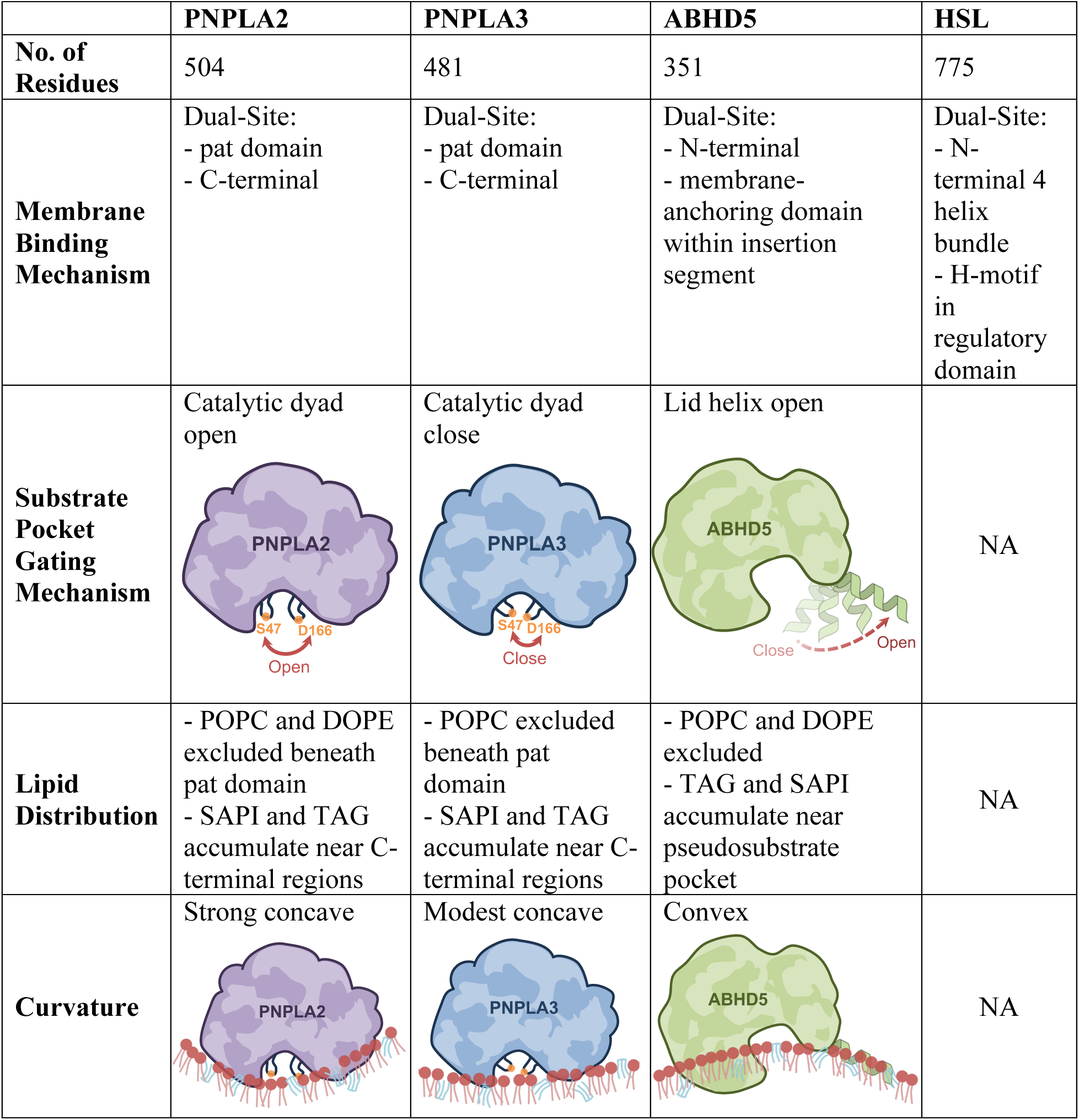
Summary of membrane-targeting mechanisms of PNPLA2, PNPLA3, and ABHD5, and HSL. Each row summarizes key characteristics of the four proteins. Figures in the third row depict the conformations of the substrate pocket gate. Figures in the last row illustrate membrane curvature beneath each protein, shown as either concave or convex. Colored protein surfaces (purple for PNPLA2, blue for PNPLA3, and green for ABHD5) include arrows indicating conformational changes in the catalytic dyad or lid helix. Phospholipids and TAG are represented in red and cyan, respectively.

|  | **PNPLA2** | **PNPLA3** | **ABHD5** | **HSL** |
| --- | --- | --- | --- | --- |
| **No. of Residues** | 504 | 481 | 351 | 775 |
| **Membrane Binding Mechanism** | Dual-Site:<br>- pat domain<br>- C-terminal | Dual-Site:<br>- pat domain<br>- C-terminal | Dual-Site:<br>- N-terminal<br>- membrane-anchoring domain within insertion segment | Dual-Site:<br>- N-terminal 4 helix bundle<br>- H-motif in regulatory domain |
| **Substrate Pocket Gating Mechanism** | Catalytic dyad open | Catalytic dyad close | Lid helix open | NA |
| **Lipid Distribution** | - POPC and DOPE excluded beneath pat domain<br>- SAPI and TAG accumulate near C-terminal regions | - POPC excluded beneath pat domain<br>- SAPI and TAG accumulate near C-terminal regions | - POPC and DOPE excluded<br>- TAG and SAPI accumulate near pseudosubstrate pocket | NA |
| **Curvature** | Strong concave | Modest concave | Convex | NA |

The C-terminal amphipathic helices serve as the primary membrane-targeting modules. In PNPLA2, helices αF-αI (residues 320-384) form the major binding interface, consistent with experimental evidence that deletion of residues 320-360 disrupts PNPLA2-LD localization. Moreover, our simulations identify residues 209-215 as membrane-interacting, aligning with experimental data showing this segment as critical for ABHD5-mediated activation. For PNPLA3, we showed that membrane binding is largely mediated by helices αH′-αK (residues 346-402), with flexible reorientation between ER and LD membranes, which agrees with experimental validation of C-terminal membrane targeting. The conformational plasticity of Arg371 corresponds to the functionally important QRLV motif (370–373), whose mutation reduces LD binding.

Amphipathic helices located at the C-termini of PNPLA2 and PNPLA3 play a central role in mediating membrane binding. These helices are characterized by alternating hydrophobic and polar residues, enabling them to interact favorably with lipid interfaces. Upon membrane contact, they undergo conformational rearrangements that enhance membrane insertion and stability. This dynamic behavior is consistent with that observed in other LD-associated proteins. For example, ABHD5 relies on helices within the insertion segment for membrane sensing and remodeling, where mutation of hydrophobic residues (Trp199, Phe210, Phe222) notably impairs LD association (52). Similarly, HSL anchors to the LD via an amphipathic helix where critical aromatic residues (Phe519, Trp526, Phe530) on its hydrophobic face are essential for binding; their mutation abolishes LD interaction (53). These findings highlight the predominant role of hydrophobic interactions, especially those involving aromatic side chains, in mediating the membrane anchoring of LD-targeting proteins.

Structural comparisons of the membrane-bound states of PNPLA2 and PNPLA3 reveal significant differences in their C-terminal architecture. In PNPLA2, αF-αI form a deeply embedded U-shaped helical bundle that engages both the membrane interface and the hydrophobic core. This anchoring conformation is associated with an open catalytic dyad geometry (Ser47-Asp166), conducive to TAG hydrolysis. PNPLA2 also penetrates deeper into the lipid layer and induces more pronounced membrane deformation than PNPLA3, indicating a more dynamic and catalytically engaged interaction with the lipid environment. In contrast, the C-terminal region of PNPLA3 is more solvent-exposed and conformationally flexible. On LDs, PNPLA3 forms a shallower inverted U-shaped helix bundle, while on ER membranes, the helices adopt more linear and extended conformations. This flexible C-terminal behavior correlates with a more compact and catalytically restrained dyad configuration, suggesting limited enzymatic activity despite membrane association.

Our simulations demonstrate that membrane binding modulates the conformation of the catalytic dyad, which serves as a gating switch for enzymatic activity. In PNPLA2, LD association promotes a transition to a fully open dyad conformation, facilitating lipid access and catalysis. PNPLA3, in contrast, samples closed or semi-open dyad states depending on membrane type, consistent with its reduced enzymatic activity (51). These adaptive structural responses underscore the role of local membrane environment in tuning catalytic function. Interestingly, this regulatory mechanism bears resemblance to the gating strategy observed in ABHD5. Although ABHD5 lacks intrinsic lipase activity, it binds LDs via a flexible lid helix that transitions between open and closed conformations, controlling access to its pseudosubstrate pocket (**Table 3**). Unlike the helical bundle-mediated anchoring in PNPLA2, ABHD5 membrane engagement is governed by conformational flexibility within the lid region, highlighting distinct structural solutions for membrane sensing and lipid gating among LD-associated proteins.

Upon binding to LDs, both PNPLA2 and PNPLA3 induce local lipid reorganization. This is characterized by the displacement of POPC and the enrichment of TAG molecules near their membrane-anchoring helices, driven by hydrophobic insertion. Notably, such lipid redistribution is not observed when proteins interact with ER membranes, indicating a distinct membrane-specific binding mode. Beyond lipid distribution changes, PNPLA2 also induces pronounced negative membrane curvature under its patatin domain This suggests an active role in LD surface remodeling that may facilitate substrate access. In contrast, PNPLA3 generates only subtle membrane perturbations. These remodeling patterns differ significantly from those of ABHD5, which produces positive curvature and forms localized TAG-enriched nanodomains beneath its pseudosubstrate pocket (**Table 3**).

In summary, our study uncovers distinct membrane-binding mechanisms for PNPLA2 and PNPLA3, governed by their C-terminal amphipathic helices and responsive to lipid environment. These structural differences influence the conformation of the catalytic dyad, revealing a dynamic gating mechanism that links membrane association to enzymatic activity. By integrating multiscale approaches with structural analysis, we provide a molecular framework for understanding how lipid membranes regulate PNPLA function. Given that PNPLA2 is essential for lipid mobilization and energy balance, while PNPLA3 variants are strongly associated with hepatic steatosis and fibrosis, our findings offer mechanistic insight into their divergent physiological roles. The ability of membrane composition to modulate catalytic accessibility suggests potential strategies for selective modulation of PNPLA activity. This work lays the foundation for rational design of mutations or small molecules that tune membrane binding or conformational plasticity, with implications for therapeutic targeting of lipid-associated metabolic diseases.

## Data, Materials, and Software Availability

The simulation input files, resulting trajectories, and post-MD analysis scripts are freely available at https://doi.org/10.5281/zenodo.18462052. All other data are included in the manuscript and/or supporting information.

## Acknowledgments

We thank the Wayne State University (WSU) High Performance Computing Center for providing the resources used for CGMD and GaMD simulations. This work was supported by WSU startup funds, WSU Postdoctoral Fellows funds, and NIH grant R35GM160192 (to Y.M.H.). Additional support was provided by NIH grant R01 DK126743 from the National Institute of Diabetes and Digestive and Kidney Diseases (to E.P.M.) and the WSU Barber Integrative Metabolic Research Program (to Y.M.H. and E.P.M.).

## Author Contributions

Conceptualization: Y.M.H. and E.P.M. Project design: A.K. and Y.M.H. Perform simulations and analyze data: A.K. and YM.H. Experimental validation: G.T. and E.P.M. Manuscript writing: A.K., G.T., E.P.M., and Y.M.H.

## Movie Legends

**Movie 1: Modeling the diffusion of PNPLA2 to the LD using CGMD.** All color representations are consistent with those in Figure 3.

**Movie 2: Modeling the diffusion of PNPLA3 to the ER using CGMD.** All color representations are consistent with those in Figure 3.

**Movie 3: Modeling the diffusion of PNPLA3 to the LD using CGMD.** All color representations are consistent with those in Figure 3.

**Movie 4: Modeling the dynamics of PNPLA2 bound to LD using all-atom GaMD.** All color representations are consistent with those in Figure 3.

**Movie 5: Modeling the dynamics of PNPLA3 bound to ER using all-atom GaMD.** All color representations are consistent with those in Figure 3.

**Movie 6: Modeling the dynamics of PNPLA3 bound to LD using all-atom GaMD.** All color representations are consistent with those in Figure 3.

