## Supplemental file for "Multiscale Analysis of PNPLA2 and PNPLA3 Membrane Targeting"

**Short Title: PNPLA2/PNPLA3 Membrane Binding**

*Amit Kumar[a], Grace Teskey[b], Emilio P. Mottillo[b,c], and Yu-ming M. Huang[a*]*

[a]Department of Physics and Astronomy, Wayne State University, Detroit, MI 48201, USA

[b]Department of Internal Medicine, Hypertension and Vascular Research Division, Henry Ford Hospital, Detroit, MI 48202, USA

[c]Department of Physiology, Wayne State University School of Medicine, Detroit, MI 48201, USA

*corresponding author:

Yu-ming M. Huang

ORCID: 0000-0003-3257-6170

| Simulation Protocol | Stage | Simulation steps/time |
|---|---|---|
| System information | $N_{beads}$ | PNPLA2 on LD: 41331<br>PNPLA3 on LD: 33806<br>PNPLA3 on ER: 40113 |
| | Box dimension | PNPLA2 on LD: 148 Å x 148 Å x 270 Å<br>PNPLA3 on LD: 147 Å x 147 Å x 265 Å<br>PNPLA3 on ER: 147 Å x 147 Å x 227 Å |
| System minimization | Minimization | 5000 Steps |
| | Minimization | 5000 steps |
| System equilibration | Equ (FC=200) | 10 ns |
| | Equ (FC=100) | 5 ns |
| | Equ (FC=50) | 2 ns |
| | Equ (FC=20) | 1 ns |
| | Equ (FC=10) | 1 ns |
| Production simulation | CGMD | 20 μs |

**Table S1:** Summary of CGMD simulation protocols. This table details the sequential steps, including the total number of beads ($N_{beads}$) per system, steepest descent minimization, and equilibration at 303 K using varied force constraints (FC: 200, 100, 50, 20, and 10 kJ/mol/Å$^2$). Production runs extended for 20 μs.

| Simulation Protocol | Stage | PNPLA2 in Sol | PNPLA2 on LD | PNPLA3 in Sol | PNPLA3 on LD | PNPLA3 on ER |
|---|---|---|---|---|---|---|
| System information | $N_{atoms}$ | 175912 | 326688 | 125518 | 327877 | 252286 |
| | Box dimension | 130Å x 109Å x 144Å | 155Å x 155Å x 190Å | 91Å x 132Å x 125Å | 153Å x 153Å x 185Å | 153Å x 153Å x 160Å |
| System minimization | $M_{Hydrogen}$ | 500 steps | - | 500 steps | - | - |
| | $M_{SideChain}$ | 5000 steps | - | 5000 steps | - | - |
| | $M_{Protein}$ | 5000 steps | - | 5000 steps | - | - |
| | $M_{Water}$ | 1000 steps | - | 1000 steps | - | - |
| | $M_{System}$ | 5000 steps | - | 5000 steps | - | - |
| | $M_{SD}$ | - | 2000 steps | - | 2000 steps | 2000 steps |
| | $M_{CG}$ | - | 3000 steps | - | 3000 steps | 3000 steps |
| System equilibration | $Equ_{0\text{-}50K}$ | 10 ps | - | 10 ps | - | - |
| | $Equ_{50\text{-}100K}$ | 10 ps | - | 10 ps | - | - |
| | $Equ_{100\text{-}150K}$ | 10 ps | - | 10 ps | - | - |
| | $Equ_{150\text{-}200K}$ | 10 ps | - | 10 ps | - | - |
| | $Equ_{200\text{-}250K}$ | 10 ps | - | 10 ps | - | - |
| | $Equ_{250\text{-}303K}$ | 10 ps | - | 10 ps | - | - |
| | $Equ_{0\text{-}100K}$ | - | 10 ps | - | 10 ps | 10 ps |
| | $Equ_{100\text{-}303K}$ | - | 200 ps | - | 200 ps | 200 ps |
| | $Equ_{Hold}$ | - | 1 ns x 10 | - | 1 ns x 10 | 1 ns x 10 |
| | cMD simulation | 20 ns | 50 ns | 20 ns | 50 ns | 50 ns |
| GaMD preparation | ntcmd | 8 ns | 12 ns | 4 ns | 12 ns | 8 ns |
| | ntebprep | 4 ns | 6 ns | 2 ns | 6 ns | 4 ns |
| | nteb | 50 ns | 60 ns | 50 ns | 60 ns | 50 ns |
| Production simulation | GaMD simulation | 1000 ns | 1000 ns | 1000 ns | 1000 ns | 1000 ns |

**Table S2:** Summary of AAMD simulation protocols. The table includes the total number of atoms ($N_{atoms}$) in each system. We used steepest descent (SD) and conjugate gradient (CG) algorithms for minimization. GaMD preparation for all systems followed three steps: 1) collecting potential energy statistics (ntcmd), 2) running GaMD with a fixed, non-updating boost potential (ntebprep), and 3) performing GaMD with an actively updated boost potential (nteb). Production GaMD simulations then used this final and fixed boost potential.

| **Primer Name** | **Primer Sequence 5'-3'** |
|---|---|
| hPNPLA2 R364E - Forward | CGAGGACATCGAGTGGATGAAGGAGC |
| hPNPLA2 R364E - Reverse | GCTCCTTCATCCACTCGATGTCCTCG |
| hPNPLA3 R349E - Forward | GCTACCCATTGAGATAATGTCTTATGTAATGC |
| hPNPLA3 R349E - Reverse | GCATTACATAAGACATTATCTCAATGGGTAGC |

**Table S3:** List of Overlap Extension PCR primers.

| Protein | Domain / Region | Polar Residues | Non-polar Residues | Positively Charged Residues | Negatively Charged Residues |
|---|---|---|---|---|---|
| **PNPLA2** | Full protein | 226 | 278 | 48 | 50 |
| | Pat domain (residue 10-180) | 73 | 98 | 14 | 13 |
| | C-terminal (residue 181-481) | 144 | 157 | 31 | 35 |
| | C-terminal (residue 482-504) | 4 | 19 | 1 | 1 |
| **PNPLA3** | Full protein | 224 | 257 | 48 | 45 |
| | Pat domain (residue 10-180) | 76 | 95 | 17 | 13 |
| | C-terminal (residue 181-481) | 143 | 158 | 27 | 33 |

**Table S4:** Summary of polar and non-polar residues in different domains of PNPLA2 and PNPLA3.

| Domain / Region | PNPLA2 on LD | PNPLA3 on LD | PNPLA3 on ER |
|---|---|---|---|
| **Overall buried residue distance** | -4.07 ± 2.78 Å | -4.83 ± 4.15 Å | -5.91 ± 3.66 Å |
| **Buried residues from pat domain** | -3.13 ± 2.44 Å | -3.07 ± 1.80 Å | -2.07 ± 1.23 Å |
| **Buried residues from C-terminal** | -4.15 ± 2.80 Å | -5.09 ± 4.34 Å | -6.40 ± 3.57 Å |
| **Deepest buried residue:**<br>P337 (αG helix) for PNP2-LD<br>C358 (αH' helix) for PNP3-LD<br>C358 (αH' helix) for PNP3-ER | -12.35 ± 1.34 Å | -15.86 ± 1.97 Å | -12.86 ± 0.69 Å |

**Table S5:** Average distance between PNPLA2 and PNPLA3 residues and membrane surface in different membrane-bound states. This table lists the average vertical distance (in Å) from each residue of PNPLA2 and PNPLA3 to the membrane surface in LD- and ER-bound systems. The measurements reflect the mean distance between the Cα atom of each residue and the phosphorus atoms of the upper membrane leaflet. Residues with negative values indicate penetration below the membrane surface. The table also summarizes the membrane penetration for key structural regions.

| System | PNPLA2 in solution | PNPLA2 on LD | PNPLA3 in solution | PNPLA3 on ER | PNPLA3 on LD |
|---|---|---|---|---|---|
| α1 | 16-33 | 19-33 | 16-33 | 17-31 | 17-31 |
| α2 | 34-39 | 34-39 | 34-40 | 34-40 | 34-40 |
| α3 | 47-59 | 48-59 | 48-59 | 48-59 | 48-59 |
| α4 | 62-78 | 62-79 | 62-77 | 64-79 | 64-79 |
| α5 | 90-102 | 93-100 | 91-102 | 91-102 | 91-102 |
| α6 | 105-111 | NA | 105-111 | 105-111 | 105-111 |
| α6’ | NA | NA | NA | 121-127 | NA |
| α7 | 135-147 | 135-145 | 135-147 | 135-147 | 135-147 |
| α8 | 167-170 | NA | 167-170 | NA | NA |
| β1 | 9-13 | NA | 10-13 | NA | 10-13 |
| β2 | 42-46 | NA | 42-47 | NA | 42-47 |
| β3 | 114-121 | NA | 114-121 | NA | 118-121 |
| β4 | 125-130 | NA | 125-130 | NA | 123-127 |
| β5 | 158-160 | NA | 158-160 | NA | NA |
| β6 | 162-166 | NA | 162-166 | NA | 164-166 |
| αA | 216-228 | 216-226 | 216-227 | 217-226 | 216-227 |
| αB | 231-253 | 231-251 | 231-253 | 234-253 | 231-253 |
| αC | 275-278 | 273-276 | 281-285 | NA | NA |
| αD | 294-298 | 294-299 | 287-299 | 287-299 | 286-299 |
| αE | 302-311 | 302-311 | 300-309 | 300-310 | 300-309 |
| αF | 318-324 | 320-325 | 314-320 | 315-320 | NA |
| αG | 326-337 | 326-337 | 322-333 | 322-333 | 325-333 |
| αH | 338-357 | 338-353 | 338-344 | NA | NA |
| αH’ | NA | NA | 346-360 | 346-360 | 346-360 |
| αI | 358-372 | 354-370 | 361-371 | 361-376 | 361-369 |
| αJ | 374-383 | 371-384 | 372-392 | 378-387 | 372-384 |
| αK | NA | NA | 394-402 | 388-401 | 388-402 |
| αL | 429-446 | 429-450 | NA | NA | NA |
| αM | NA | NA | 450-459 | NA | NA |
| βA | 181-184 | NA | 181-184 | NA | 181-184 |

**Table S6:** Summary of secondary structure elements of PNPLA2 and PNPLA3 in different environments. The table lists the secondary structure identified from the modeling of PNPLA2 and PNPLA3 in solution, LD-bound, and ER-bound states. Helices α1-α8 and strands β1-β6 correspond to the patatin domain, while helices αA-αM and strand βA represent the C-terminal region. “NA” indicates that no stable secondary structure was identified for given sequence.

| Domain / Region | PNPLA2 in solution | PNPLA2 on LD | PNPLA3 in solution | PNPLA3 on ER | PNPLA3 on LD |
|---|---|---|---|---|---|
| **Patatin domain:**<br>residues 10-180 | 0.85 ± 0.17 | 3.18 ± 0.32 | 0.84 ± 0.21 | 2.62 ± 0.59 | 3.40 ± 0.51 |
| **C-terminal:**<br>residues 181-504 for PNP2<br>residues 181-481 for PNP3 | 1.98 ± 1.25 | 4.31 ± 1.31 | 1.57 ± 0.82 | 3.55 ± 1.10 | 4.92 ± 1.42 |
| **Loop/helix in C-terminal:**<br>residues 181-325 for PNP2-sol<br>residues 181-325 for PNP2-LD<br>residues 181-321 for PNP3-sol<br>residues 181-321 for PNP3-ER<br>residues 181-324 for PNP3-LD | 1.74 ± 1.57 | 3.71 ± 0.85 | 1.34 ± 0.81 | 3.34 ± 1.28 | 4.22 ± 0.86 |
| **αF in C-terminal** | 1.64 ± 0.14 | 3.64 ± 0.10 | 0.87 ± 0.18 | 3.23 ± 0.22 | NA |
| **αG in C-terminal** | 2.00 ± 0.21 | 3.70 ± 1.30 | 0.76 ± 0.13 | 2.86 ± 0.24 | 4.06 ± 0.20 |
| **αH in C-terminal** | 1.59 ± 0.18 | 3.32 ± 0.11 | 2.28 ± 0.32 | NA | NA |
| **αH' in C-terminal** | NA | NA | 1.68 ± 0.22 | 3.50 ± 0.31 | 4.58 ± 0.44 |
| **αI in C-terminal** | 1.14 ± 0.21 | 3.21 ± 0.10 | 1.32 ± 0.18 | 3.06 ± 0.15 | 3.78 ± 0.10 |
| **αJ in C-terminal** | 2.35 ± 0.60 | 3.24 ± 0.21 | 1.25 ± 0.22 | 3.66 ± 0.29 | 4.69 ± 0.61 |
| **αK in C-terminal** | NA | NA | 1.12 ± 0.07 | 3.98 ± 0.10 | 5.55 ± 0.64 |
| **αL in C-terminal** | 1.37 ± 0.41 | 4.89 ± 0.27 | NA | NA | NA |
| **Terminal loop region:**<br>residues 447-504 for PNP2-sol<br>residues 451-504 for PNP2-LD<br>residues 403-481 for PNP3-sol<br>residues 402-481 for PNP3-ER<br>residues 403-481 for PNP3-LD | 2.16 ± 0.58 | 6.33 ± 1.33 | 2.16 ± 0.83 | 3.96 ± 1.08 | 6.39 ± 1.62 |

**Table S7:** RMSF analysis of key structural regions of PNPLA2 and PNPLA3 across different environments. The table summarizes the average RMSF values (Å) calculated from GaMD simulations for selected structural regions of PNPLA2 and PNPLA3. Refer to Table S6 for the corresponding residue numbers for each helix (αF-αL).

| Domain / Region | PNPLA2 in solution | PNPLA2 on LD | PNPLA3 in solution | PNPLA3 on ER | PNPLA3 on LD |
|---|---|---|---|---|---|
| **Rg** | 25.52 ± 0.79 | 36.85 ± 0.53 | 23.92 ± 0.63 | 27.08 ± 0.20 | 26.52 ± 0.17 |
| **No. of H-bonds** | 239 ± 7 | 205 ± 8 | 227 ± 7 | 197 ± 7 | 194 ± 7 |

**Table S8:** Radius of gyration (Rg) and intraprotein hydrogen bond analysis of PNPLA2 and PNPLA3 in different environments. This table summarizes the average Rg values (Å) and the number of intraprotein hydrogen bonds calculated from GaMD simulations of PNPLA2 and PNPLA3 in solution, LD-bound, and ER-bound states. Values represent the mean across three simulation replicas, which reflects changes in protein compactness and internal stabilization upon membrane association.

| Lipid Components | PNPLA2 on LD | PNPLA3 on LD | PNPLA3 on ER |
|---|---|---|---|
| **POPC** (residues)<br>(atoms) | 4.58 ± 0.68<br>112.62 ± 39.56 | 3.06 ± 0.57<br>92.00 ± 14.56 | 0.44 ± 0.38<br>1.21 ± 1.60 |
| **DOPE** (residues)<br>(atoms) | 0.21 ± 0.29<br>2.26 ± 3.36 | 0.67 ± 0.44<br>5.77 ± 5.98 | 1.12 ± 0.18<br>15.31 ± 7.96 |
| **SAPI** (residues)<br>(atoms) | 0.95 ± 0.11<br>29.53 ± 15.33 | 0.03 ± 0.05<br>0.13 ± 0.22 | 2.00 ± 0.00<br>55.19 ± 8.67 |
| **TAG** (residues)<br>(atoms) | 2.00 ± 1.08<br>44.85 ± 29.85 | 0.51 ± 0.52<br>6.72 ± 9.69 | NA |
| **Total PL** (residue)<br>(atom) | 5.74 ± 1.35<br>144.41 ± 33.19 | 3.76 ± 0.92<br>97.90 ± 29.73 | 3.56 ± 0.45<br>71.71 ± 16.16 |
| **Total** (residues)<br>(atoms) | 7.74 ± 0.95<br>189.26 ± 23.48 | 4.27 ± 0.68<br>104.62 ± 21.99 | 3.56 ± 0.45<br>71.71 ± 16.16 |

**Table S9:** Average number of lipid molecules within 10 Å of the catalytic dyad residues Ser47 and Asp166 in PNPLA2 and PNPLA3 across different membrane-bound states. Values represent means ± standard deviations calculated from three independent GaMD simulation replicas.

A) PNPLA2 in solution

Replica 1

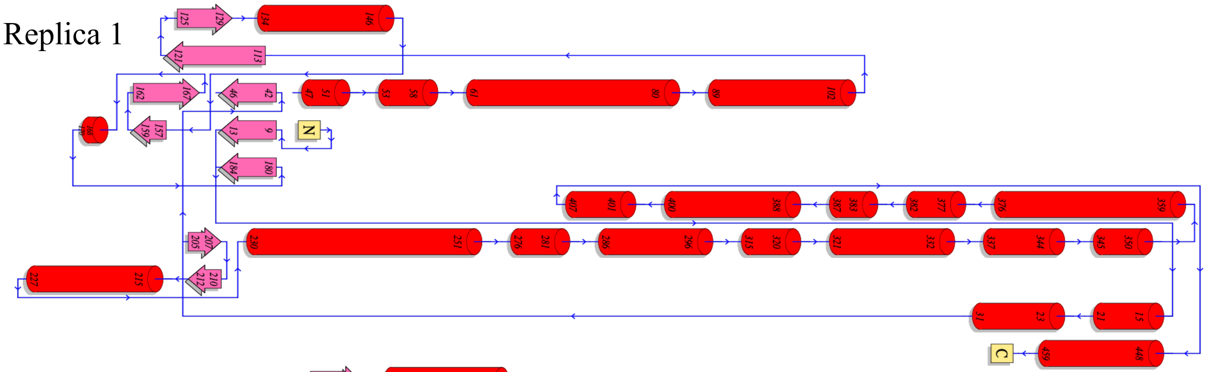


Replica 2

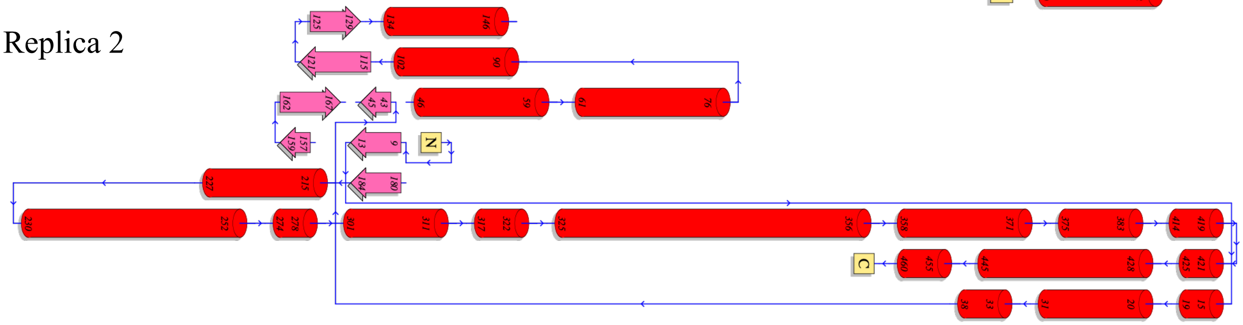


Replica 3

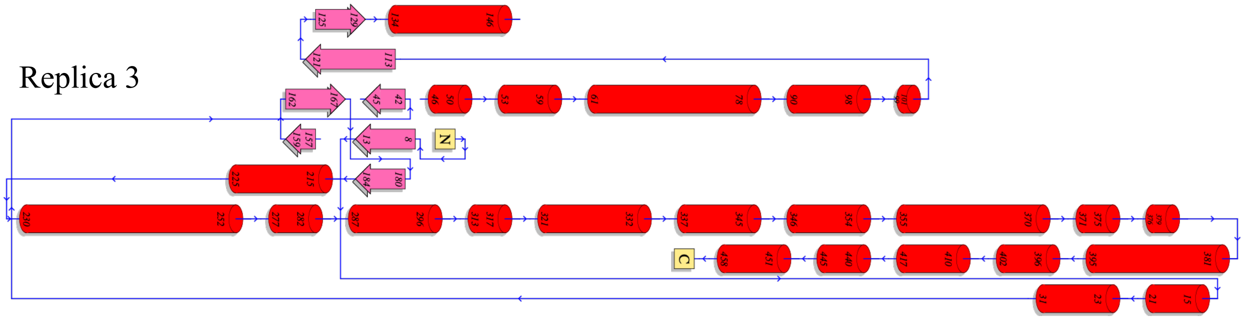


(B) PNPLA3 in solution

Replica 1

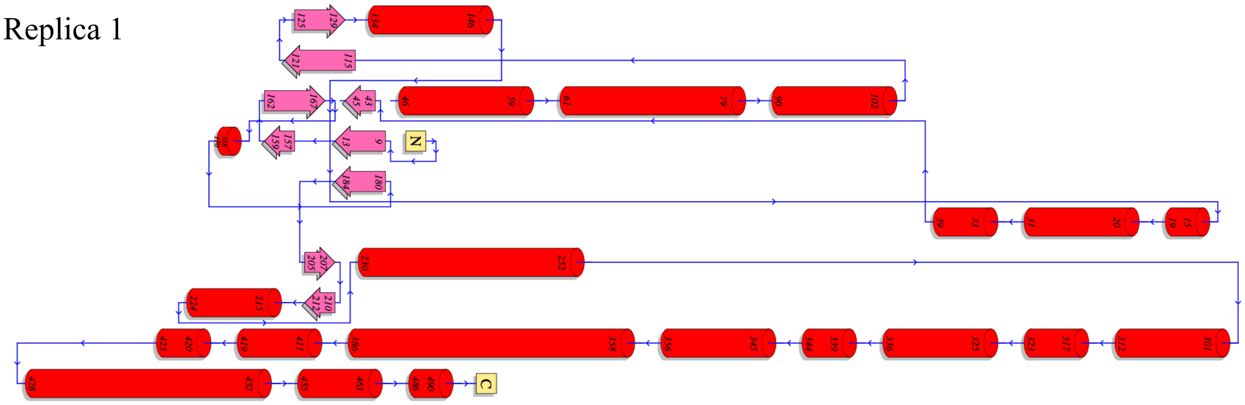

Replica 2

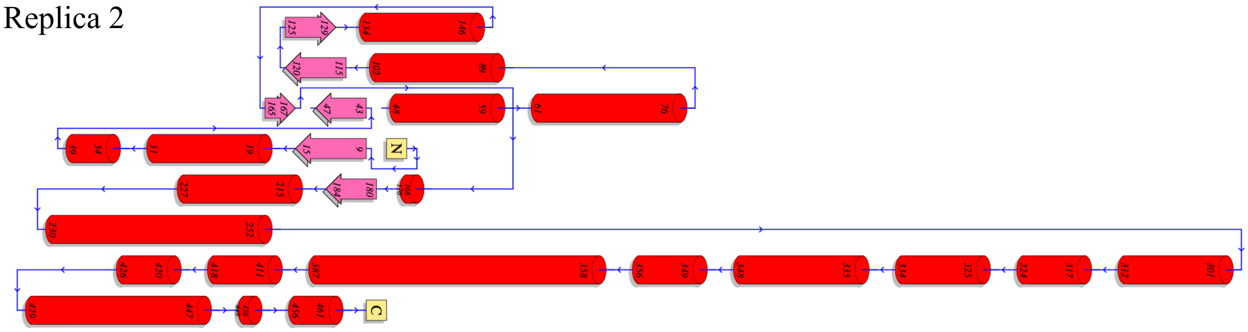

Replica 3

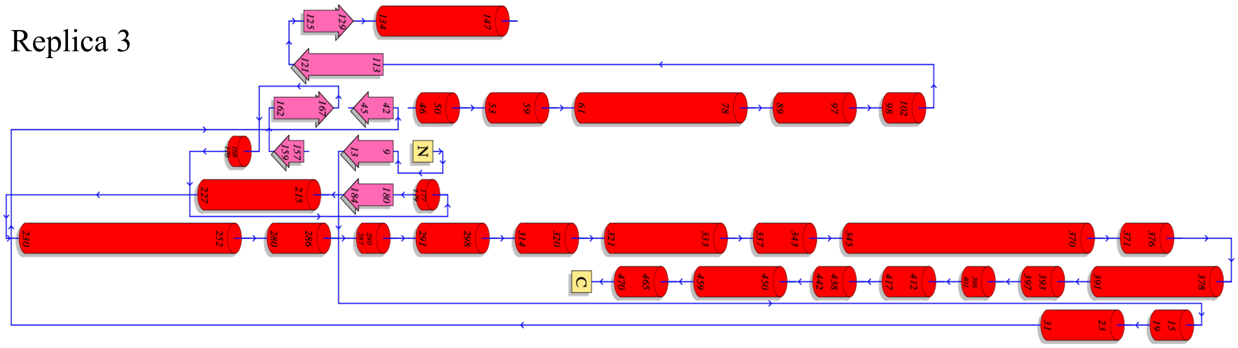

(C) PNPLA2 on LD

Replica 1

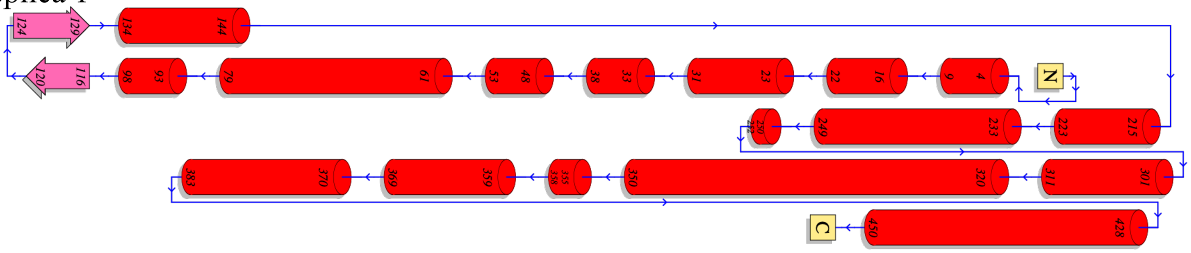

Replica 2

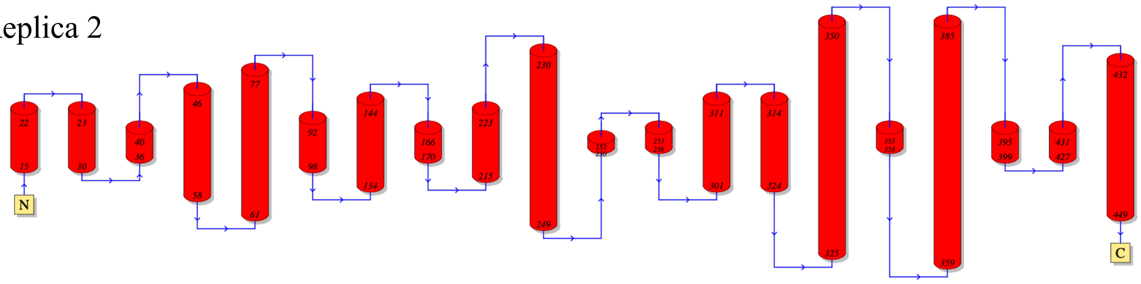

Replica 3

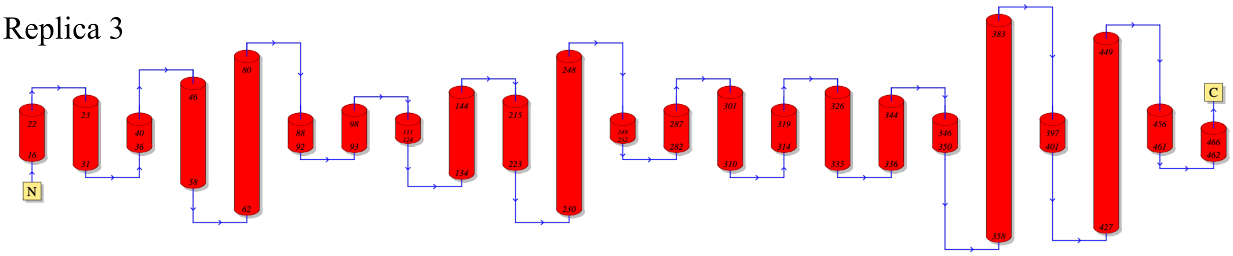

(D) PNPLA3 on ER

Replica 1

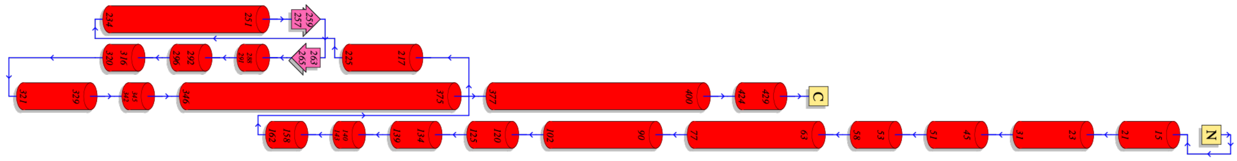


Replica 2

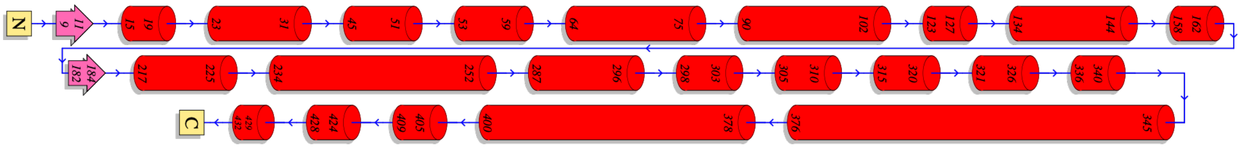


Replica 3

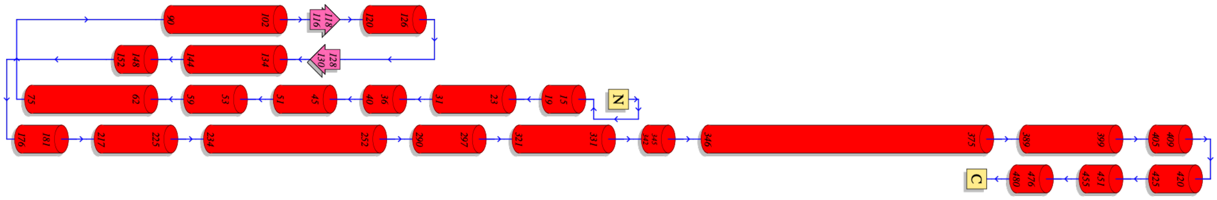


(E) PNPLA3 on LD

Replica 1

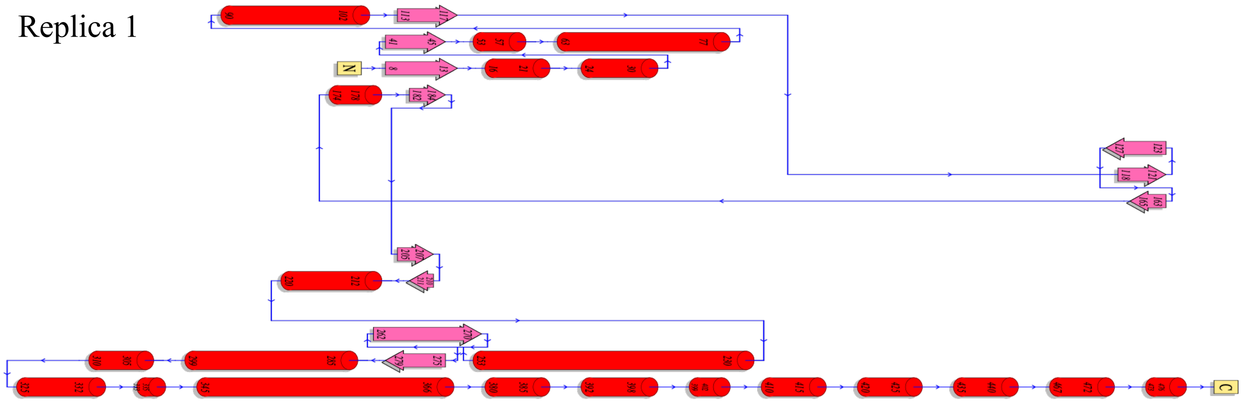


Replica 2

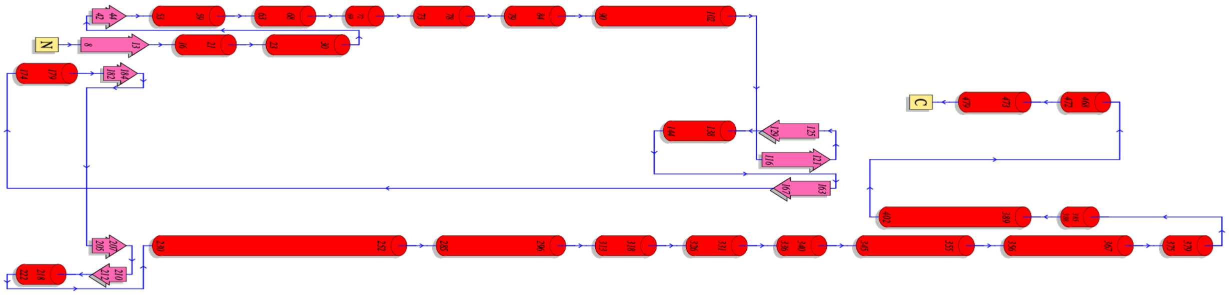

Replica 3

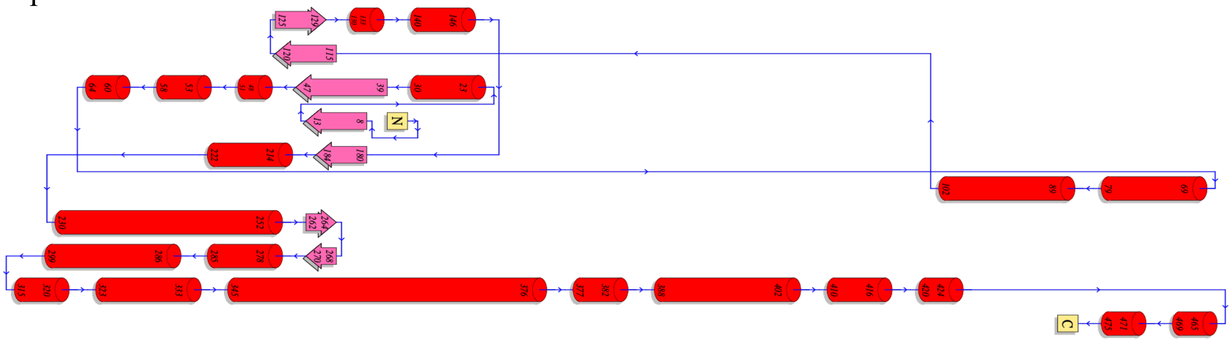


**Figure S1:** Secondary structure analysis of PNPLA2 and PNPLA3 in solution, ER-bound, and LD-bound states across three replica MD simulations.

| | PNPLA2 | PNPLA3 |
|---|---|---|
| Solution | 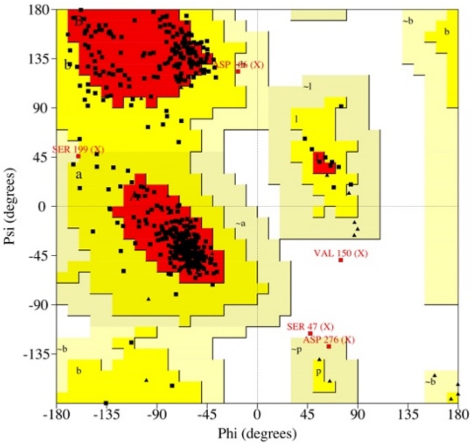  | 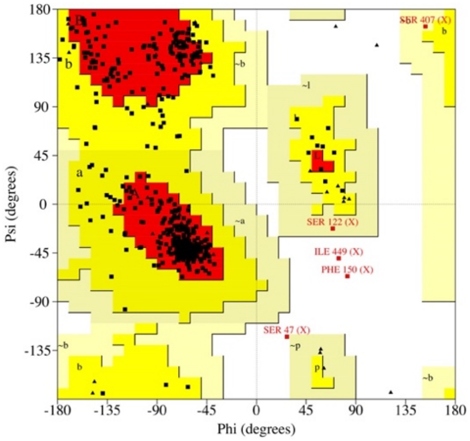  |
| LD | 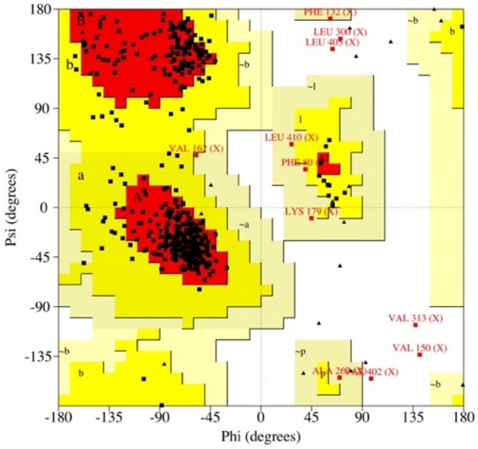  | 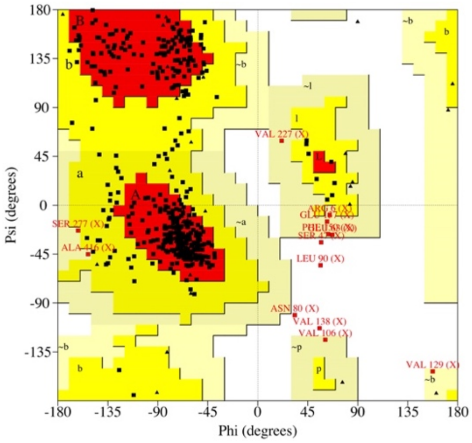  |
| ER | NA | 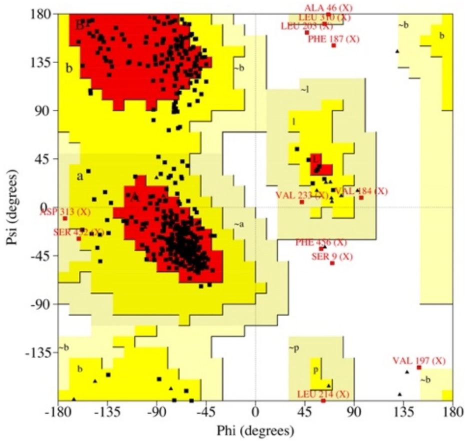  |

**Figure S2:** Ramachandran plot analysis of representative MD replicas of PNPLA2 and PNPLA3. The backbone dihedral angles (ϕ and ψ) were evaluated from MD simulations of PNPLA2 and

PNPLA3 in three environments: solution, LD-bound, and ER-bound. Each plot displays the distribution of residues in favored, allowed, and disallowed regions. High percentages of residues in the favored regions support the structural quality and reliability of the modeled proteins under each condition.

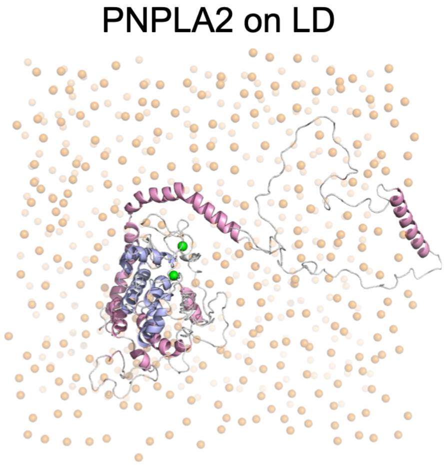


**Figure S3:** Representative top-view snapshots of membrane-bound systems from the simulations. Lipid headgroup positions are shown as orange spheres to depict the membrane surface. The protein is shown as cartoon with the patatin domain in pink and the C-terminal domain in light blue. The catalytic residues Ser47 and Asp166 are shown as green spheres.

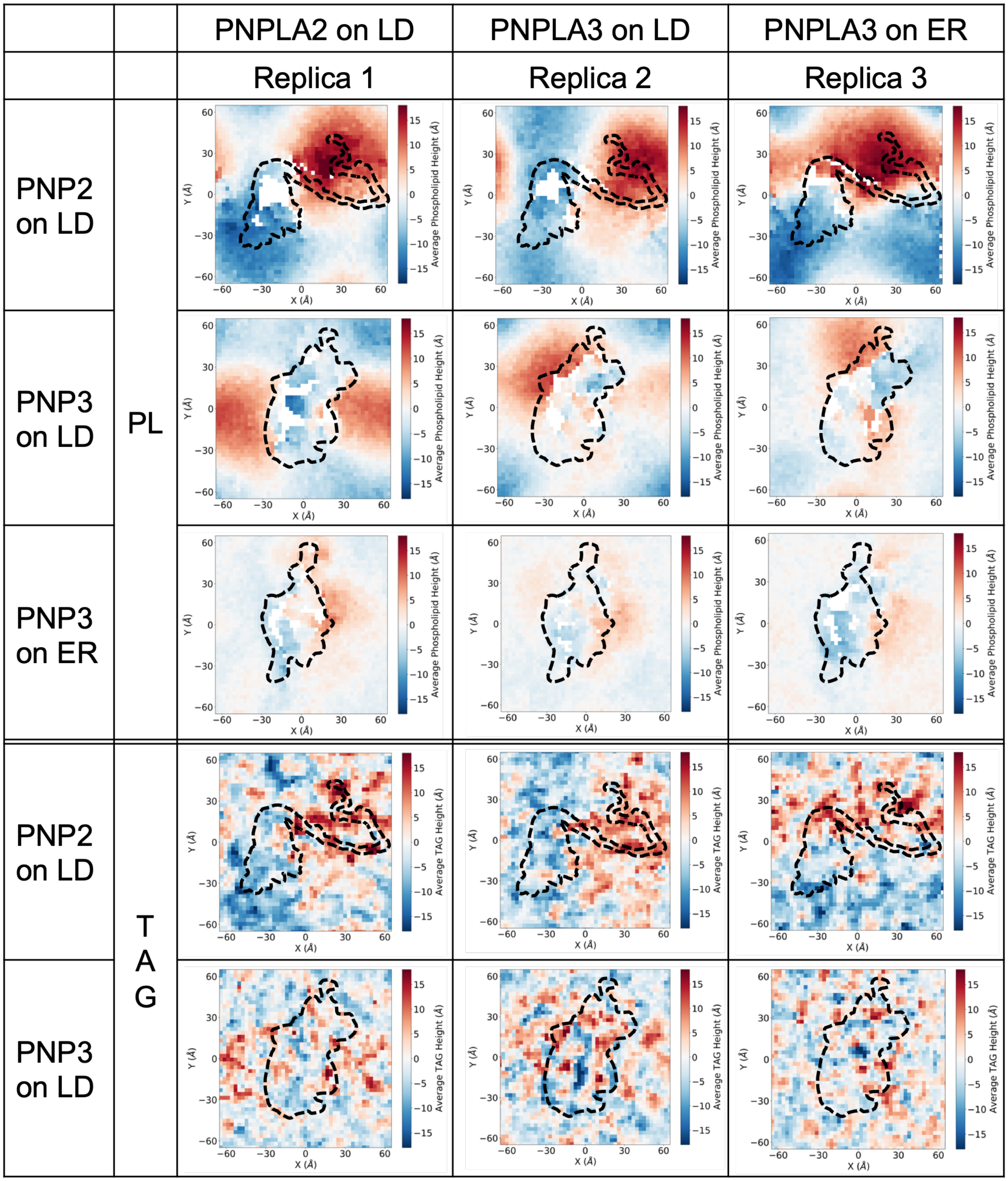


**Figure S4:** Membrane curvature induced by PNPLA2 and PNPLA3 binding. This figure supplements Figure 8 by showing results from all three replicates for each system.
